# LIGHTHOUSE illuminates therapeutics for a variety of diseases including COVID-19

**DOI:** 10.1101/2021.09.25.461785

**Authors:** Hideyuki Shimizu, Manabu Kodama, Masaki Matsumoto, Yasuko Orba, Michihito Sasaki, Akihiko Sato, Hirofumi Sawa, Keiichi I. Nakayama

**Affiliations:** Department of Molecular and Cellular Biology, Medical Institute of Bioregulation, Kyushu University, Fukuoka 812-8582, Japan; Department of Systems Biology, Harvard Medical School, Boston, MA 02115, USA; Wyss Institute for Biologically Inspired Engineering, Harvard Medical School, Boston, MA 02115, USA; Department of AI Systems Medicine, M&D Data Science Center, Tokyo Medical and Dental University, Tokyo 113-8510, Japan; Department of Omics and Systems Biology, Niigata University Graduate School of Medical and Dental Sciences, Niigata 951-8510, Japan; Division of Molecular Pathobiology, International Institute for Zoonosis Control, Hokkaido University, Sapporo 060-8638, Japan; Drug Discovery and Disease Research Laboratory, Shionogi & Co. Ltd., Osaka 561-0825, Japan; International Collaboration Unit, International Institute for Zoonosis Control, Hokkaido University, Sapporo 060-8638, Japan; One Health Research Center, Hokkaido University, Sapporo 060-8638, Japan; Global Virus Network, Baltimore, MD 21201, USA

**Keywords:** Artificial intelligence, drug discovery, drug repurposing, COVID-19

## Abstract

One of the bottlenecks in the application of basic research findings to patients is the enormous cost, time, and effort required for high-throughput screening of potential drugs for given therapeutic targets. Here we have developed LIGHTHOUSE, a graph-based deep learning approach for discovery of the hidden principles underlying the association of small-molecule compounds with target proteins. Without any 3D structural information for proteins or chemicals, LIGHTHOUSE estimates protein-compound scores that incorporate known evolutionary relations and available experimental data. It identified novel therapeutics for cancer, lifestyle-related disease, and bacterial infection. Moreover, LIGHTHOUSE predicted ethoxzolamide as a therapeutic for coronavirus disease 2019 (COVID-19), and this agent was indeed effective against alpha, beta, gamma, and delta variants of severe acute respiratory syndrome coronavirus 2 (SARS-CoV-2) that are rampant worldwide. We envision that LIGHTHOUSE will bring about a paradigm shift in translational medicine, providing a bridge from bench side to bedside.

## Introduction

Despite enormous efforts to eradicate serious medical conditions such as cancer and infectious diseases, the translation of innovative research results into clinical practice progresses slowly,^1^ leaving a large gap between bench side and bedside. The difficulty in identifying bioactive chemicals for a given target protein is one reason for this slow progress, with high-throughput screening (HTS) of a sufficiently diverse compound library being required for each target. About 10^60^ natural compounds with a molecular mass of <500 Da are thought to exist,^2^ but HTS in most cases has been performed with only ∼10^6^ compounds. Over the past few decades, molecular docking simulations have become widely adopted to reduce the cost, time, and effort required for HTS. This approach has been successful for some proteins whose crystal structures have been solved. More recently, with the advent of AlphaFold2,^3^ the ability to predict protein structures has been greatly improved, but it remains difficult to identify pockets of proteins that are potential drug targets and drug discovery without three-dimensional (3D) structural information therefore remains a challenge. Given that high-resolution 3D structural data are not available for most proteins to date and the high computational requirements of molecular docking simulations, the application of this approach has been limited.

Recent advances in artificial intelligence (AI) have demonstrated its potential in the pharmaceutical industry.^4^ Although many AI-based drug discovery methods have been proposed, they have had limited success in translational medicine. Whereas some studies have presented AI models with biological validation experiments,^5^ many others have performed only computer-based validation without proof-of-concept biomedical experiments.^6–8^ In addition, most platforms to date have been trained with small data sets, such as Directory of Useful Decoys Enhanced (DUD-E), that have known biases^9^ and are far from reflecting real-world data. Furthermore, many existing methods are based on a single network structure, whereas ensemble learning, which combines multiple network structures with different properties, might be expected to be more accurate and appropriate for AI-based drug discovery.^10^ As far as we are aware, no published study has described the discovery and validation of therapeutics for multiple human diseases based on the use of a single AI platform.

With this background, we have developed a new AI-based drug discovery platform, designated LIGHTHOUSE (<u>L</u>ead Identification with a <u>G</u>rap<u>H</u>-ensemble network for arbitrary <u>T</u>argets by <u>H</u>arnessing <u>O</u>nly <u>U</u>nderlying primary <u>SE</u>quence), an ensemble, end-to-end, graph-based deep learning tool that can predict chemicals able to interact with any protein of interest without 3D structural information. We have applied LIGHTHOUSE to malignant, infectious, and metabolic diseases. In addition, we show that LIGHTHOUSE successfully discovered a drug effective against wild-type and variant forms of severe acute respiratory syndrome coronavirus 2 (SARS-CoV-2), with this drug already having been approved for other purposes. We therefore believe that LIGHTHOUSE will promote drug discovery by identifying, from the vast chemical space, candidate compounds for a given protein with a reduced cost, time, and effort and with a wide range of potential biomedical applications.

## Results

### LIGHTHOUSE predicts confidence and IC_50_-related scores for any protein-chemical pair

We developed an end-to-end framework that relies on a message passing neural network (MPNN) for compound embedding^11^ to calculate scores for the association between any protein and any chemical. This chemical encoder takes simplified molecular-input line-entry system (SMILES) chemical encoding as input, considers the compounds as (mathematical) graph structures, and transforms them into low-dimensional vector representations. We adopted three different embedding methods for protein sequences: CNN (convolutional neural network),^6^ Transformer,^12^ and AAC (amino acid composition up to 3-mers).^13^ These methods take amino acid sequences and embed them in numerical vectors that take into account nearby (CNN) or distant (Transformer) sequences or physicochemical properties (AAC). The products of these chemical and protein encoding steps are then concatenated and entered into a feed-forward dense neural decoder network. Each chemical-protein pair is converted into a single score after this series of computations (Fig. 1A). We used this architecture to estimate both the confidence level for chemical-protein pairs and their median inhibitory concentration (IC_50_) values.

**Figure 1.**
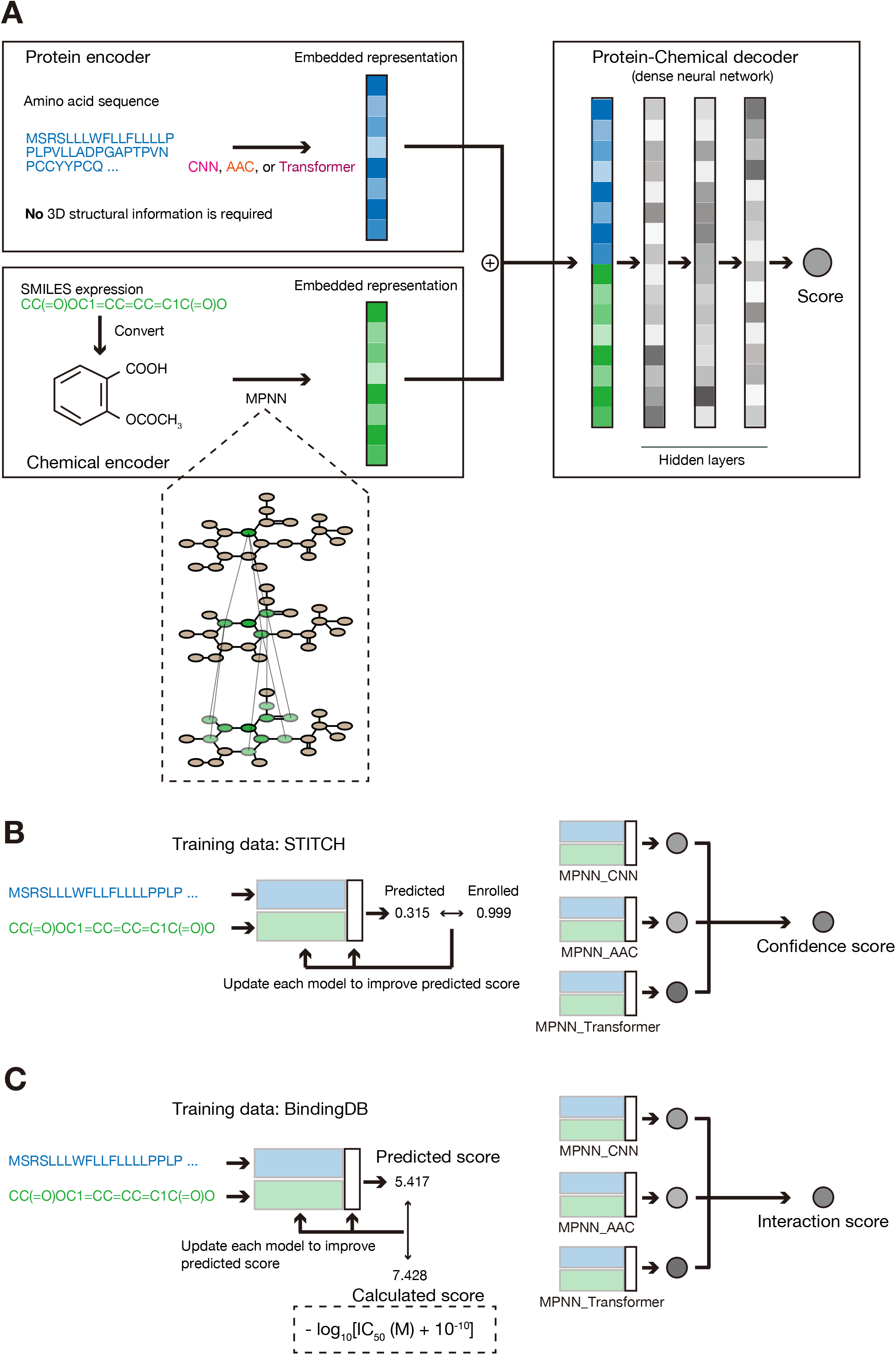
Development of LIGHTHOUSE for discovery of drug candidates without 3D structural data. (A) The basic network structure of LIGHTHOUSE consists of encoder and decoder networks. The basic network encodes the amino acid sequence of the protein of interest as numerical vectors by one of three independent methods: CNN, AAC, and Transformer. It also takes the SMILES representation of each small-molecule compound and computes the neural representation with the MPNN algorithm. The network then concatenates the protein and compound representations and calculates a “Score.” (B and C) LIGHTHOUSE consists of two modules. Module 1 estimates the association between a given compound-protein pair, and module 2 predicts a scaled IC_50_ value for the pair. In each module, the three different streams of the basic network (MPNN_CNN, MPNN_AAC, and MPNN_Transformer) are used, and the harmonic mean of the three scores is presented as the final ensemble score. Each of the three streams in module 1 (B) is trained to minimize the error between the predicted “Score” and the score registered in the STITCH database, which contains millions of known and estimated CPIs. The higher the confidence score (closer to 1), the more confident LIGHTHOUSE is that there is some relation between the compound and the protein; conversely, the lower the confidence score (closer to 0), the more confident LIGHTHOUSE is that there is no such relation. Each of the three streams in module 2 (C) is trained to predict scaled IC_50_ values with the use of BindingDB data. For instance, an interaction score of 4 means that, if the compound has inhibitory activity, the IC_50_ would be ∼100 μM, whereas an interaction score of 9 means that, if the compound has inhibitory activity, the IC_50_ would be ∼1 nM. Note that module 2 only works if the compound and protein interact, so this module is auxiliary to module 1. See also Supplementary Figs. 1,2, and Supplementary Tables 1, 2, 3.

To train the platform to estimate confidence, we used ∼1.3 million compound–(human) protein interactions (CPIs) stratify-sampled from STITCH (Table S1), which is one of the largest CPI databases^14^ and registers compound-protein pairs together with confidence scores. These scores are based on experimental data, evolutionary evidence such as homologous protein and compound relations, and co-occurrence frequencies in literature abstracts (scores range from 0 to 1, with 1 being the most reliable). To avoid overfitting, we randomly divided the overall data into training (80%), validation (10%), and test (10%) data sets (Supplementary Fig. 1A).

We fed the network with protein primary structures and chemicals and trained it to output the scores from the STITCH training data set (Fig. 1B). When we trained the three models (CNN, AAC, and Transformer for protein encoders) separately, the mean squared error (MSE) for the validation data was gradually decreased, and the area under the receiver operating characteristic curve (AUROC) was also improved (Supplementary Fig. 1B–1G). These findings indicated that our AI models learned the approximation of the hidden 1D relation underlying the compound-protein pairs without overfitting the training data. We examined the performance of the models with the test data set at the end of the training and (epoch-wise) validation phases, and we discovered that the AUROC for all three models was >0.80 (Supplementary Table 2). These scores are equivalent to or better than those of cutting-edge 3D docking simulations.^15–17^ It is of note that our AI models can be applied to proteins for which 3D structural information is not available. We took the harmonic mean of the three scores to define the confidence score (Fig. 1B).

We also trained the models to predict scores based on IC_50_ values. For this purpose, we used data from BindingDB,^18^ which collects a variety of experimental findings, and we divided the data into training (80%), validation (10%), and test (10%) data sets (Supplementary Fig. 2A). The same architecture was adopted to train the AI models to predict scaled IC_50_ values (Fig. 1C), yielding an interaction score, and we confirmed that the models adequately learned how to predict IC_50_ from amino acid sequence–chemical pairs (Supplementary Fig. 2B–2G). Finally, we assessed the performance of the models with undisclosed test data, finding that they performed well in predicting IC_50_ (Supplementary Table 3).

### LIGHTHOUSE architecture outperforms state-of-the-art methods

We next compared the performance of LIGHTHOUSE with that of similar existing methods. To ensure a fair comparison, we used DUD-E data as were used in previous studies.^7,19^ In brief, we randomly split DUD-E data (102 target proteins) into training (72 proteins) and test (30 proteins) data and then trained the LIGHTHOUSE architecture with the DUD-E training data for classification of compounds as active or decoy with regard to the protein in question. After this training, we examined LIGHTHOUSE performance with the DUD-E test data (Supplementary Fig. 3A). Of note, we used only amino acid sequences of the proteins for training and evaluation of LIGHTHOUSE, even though DUD-E provides structural data (as PDB files) for proteins. In addition, we used the balanced data set of DUD-E—the training samples comprise 22,886 active (positive) and 22,886 decoy (negative) samples—as in a previous study.^7^ LIGHTHOUSE yielded an AUROC for the DUD-E test data of 0.956 (Supplementary Fig. 3B), which was higher than the values produced by state-of-the-art methods including 3D-CNN,^20^ AtomNet,^19^ and a graph-based deep learning method proposed by Tsubaki et al.^7^ (Supplementary Fig. 3C).

For further comparison of LIGHTHOUSE with the second best method, we downloaded CPI data for human and *Caenorhabditis elegans* from the Github repository of Tsubaki et al.^7^ Both of these data sets were generated previously.^21^ Similar to Tsubaki et al., we retrained the LIGHTHOUSE architecture with these training data, and we found that LIGHTHOUSE outperformed this other method on the basis of both AUROC and F1 metrics (Supplementary Table 4). These bodies of evidence thus show that LIGHTHOUSE is one of the best architectures for drug discovery available to date.

### *In silico* verification of LIGHTHOUSE

We next evaluated the performance of LIGHTHOUSE in terms of its ability to predict known CPIs. We generated two data sets for this purpose: a “Positive” data set consisting of reliable CPIs (STITCH confidence score of >0.9), and a “Negative” data set in which the amino acid sequences of the “Positive” data set were inverted so that they would no longer be expected to interact with the corresponding chemicals. Calculation by LIGHTHOUSE of the confidence scores for both data sets revealed that those for the “Positive” data set were heavily skewed toward 1 (Figure 2A). Receiver operating characteristic (ROC) curve analysis showed that the two data sets could be distinguished on the basis of their LIGHTHOUSE confidence scores (Figure 2B). Given that the STITCH database used for the training of LIGHTHOUSE relies not only on experimental CPI data but also on co-appearance of chemicals and proteins in the literature, some well-studied molecules, such as ATP, have high values even in the “Negative” data set. Despite the presence of such false positives, LIGHTHOUSE proved to be effective in predicting the degree of association between protein-chemical pairs solely on the basis of protein primary structure.

**Figure 2.**
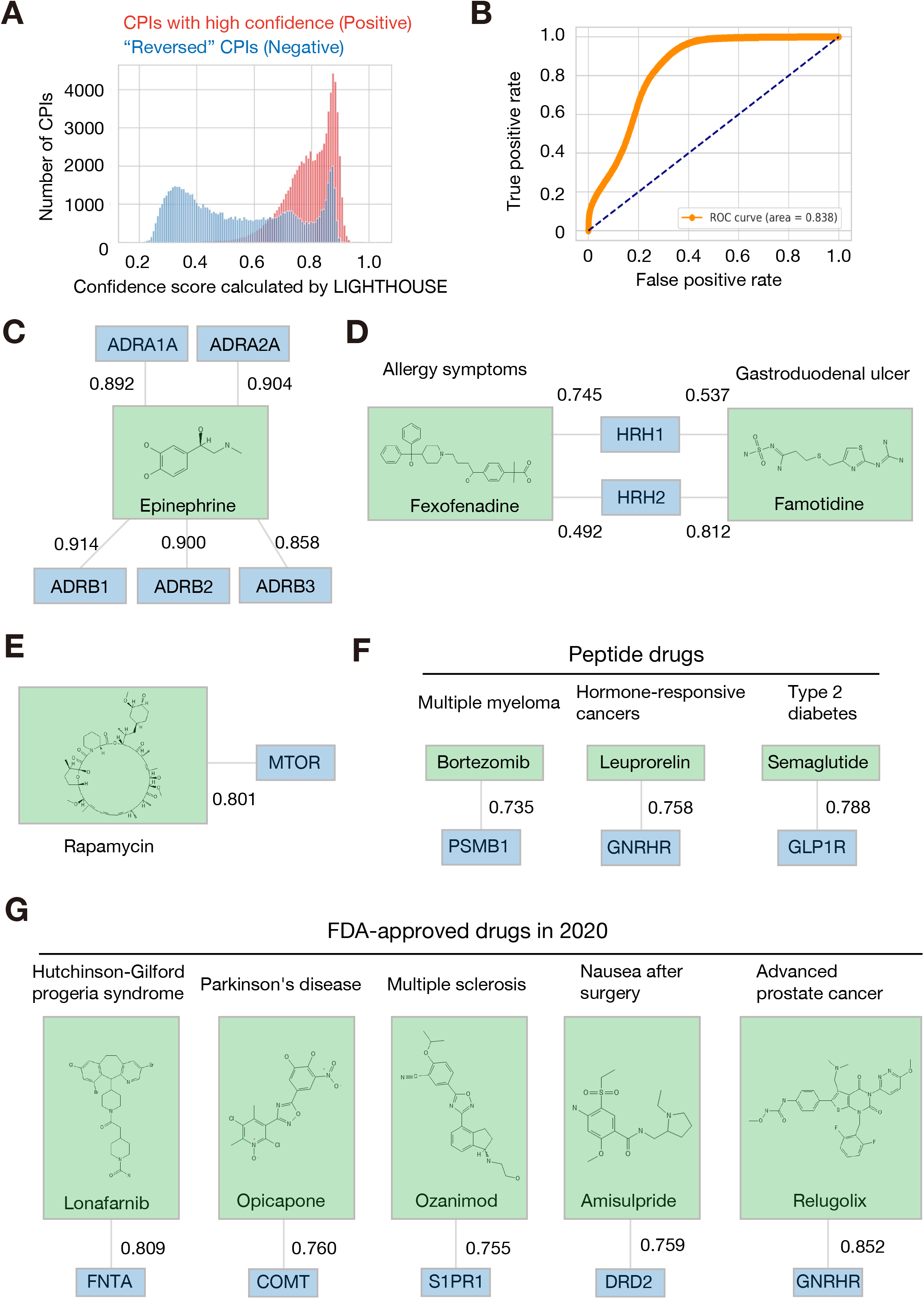
*In silico* verification of LIGHTHOUSE. (A) For investigation of whether LIGHTHOUSE is able to enrich for compounds with known targets, two data sets were generated from STITCH: a “Positive” data set consisting of CPIs with high scores (>0.9), and a “Negative” data set consisting of the same CPIs but with the amino acid sequences of the proteins reversed (for example, MTSAVM to MVASTM). Proteins in the “Negative” data set would not be expected to interact with the corresponding compounds. LIGHTHOUSE tended to yield higher confidence scores for CPIs in the “Positive” data set, with the exception of the rightmost peak for the “Negative” data set, presumably because these chemicals (such as ATP) are well known and frequently mentioned in the PubMed literature. (B) ROC curve showing that LIGHTHOUSE was able to distinguish the “Positive” and “Negative” data sets. (C–F) Known CPIs and their confidence scores predicted by LIGHTHOUSE. (C) Epinephrine and α-adrenergic (ADRA) and β-adrenergic (ADRB) receptors. (D) Fexofenadine and the histamine receptor HRH1, and famotidine and the histamine receptor HRH2. (E) The macrocyclic drug rapamycin and MTOR. (F) The peptide drugs bortezomib, leuprorelin, and semaglutide and their targets PSMB1, GNRHR, and GLP1R, respectively. These peptide drugs were converted to numerical vectors with the use of SMILES expression and MPNN. (G) Application of LIGHTHOUSE to five drugs approved by the FDA in 2020 that were not included in the training data set (published in 2016). FNTA, protein farnesyltransferase/geranylgeranyltransferase type–1 subunit α; COMT, catechol O-methyltransferase; S1PR1, sphingosine 1-phosphate receptor 1; DRD2, D2 dopamine receptor. See also Supplementary Fig. 3.

We next validated the effectiveness of LIGHTHOUSE for well-studied compound-protein pairs. LIGHTHOUSE yielded high confidence scores for adrenergic receptors (α1, α2, β1, β2, and β3) and epinephrine (Fig. 2C). Histamine receptors are classified into four subtypes,^22^ with HRH1 and HRH2 being targets of antiallergy and antiulcer drugs, respectively. LIGHTHOUSE predicted that the HRH1 antagonist fexofenadine would associate to a greater extent with HRH1 than with HRH2, whereas the HRH2 inhibitor famotidine would associate to a greater extent with HRH2 than with HRH1 (Fig. 2D). These results suggested that LIGHTHOUSE is able to accurately discriminate receptor subtype–level differences solely on the basis of amino acid sequences.

LIGHTHOUSE also proved informative both for macrocyclic chemicals such as rapamycin, yielding a high confidence score for this drug and mechanistic target of rapamycin (MTOR) (Fig. 2E), as well as for peptide drugs such as bortezomib (used for treatment of multiple myeloma), leuprorelin (hormone-responsive cancers), and semaglutide (type 2 diabetes) (Fig. 2F), yielding high confidence scores for these drugs and their known targets: proteasome subunit PSMB1,^23^ gonadotropin-releasing hormone receptor (GNRHR),^24^ and glucagon-like peptide–1 (GLP-1) receptor (GLP1R),^25^ respectively. Given the rapidly growing demand for peptide drugs,^26^ LIGHTHOUSE will prove useful for the development of novel peptide therapeutics for a variety of promising targets.

We also applied LIGHTHOUSE to five drugs that were approved by the U.S. Food and Drug Administration (FDA) in 2020 but which had not yet been registered in the STITCH database. LIGHTHOUSE successfully predicted the association between these new drugs and their target proteins (Fig. 2G), indicating the expandability of LIGHTHOUSE to a much larger exploration space than that encompassed by STITCH.

Furthermore, we also evaluated IC_50_ data from BindingDB, which is derived from actual bioassays. We found that the predicted value and observed value correlate well in the BindingDB test data set (Supplementary Fig. 4). This series of findings thus demonstrated the ability of LIGHTHOUSE to discover new drugs for a broad spectrum of diseases.

### LIGHTHOUSE discovers an inhibitor of PPAT, a key metabolic enzyme for cancer treatment

We investigated whether LIGHTHOUSE can identify compounds for novel and potentially important therapeutic targets. As such a target, we chose phosphoribosyl pyrophosphate amidotransferase (PPAT), a rate-limiting enzyme in the *de novo* nucleotide synthesis pathway, given that its expression is most correlated among all metabolic enzymes with poor prognosis in various human cancers and that its depletion markedly inhibits tumor growth^27^. Although no PPAT inhibitor has been developed and the 3D structure of the protein has not been solved, we attempted to discover an inhibitor for PPAT by LIGHTHOUSE solely on the basis of its amino acid sequence. We virtually screened ∼10^9^ commercially available compounds in the ZINC database^28^ (Supplementary Fig. 5). To reduce the calculation time, we adopted a step-by-step application of LIGHTHOUSE (Fig. 3A). The MPNN_CNN model excluded most of the chemicals unrelated to PPAT, with only 2.41% of the starting compounds having a score of >0.5 in this initial screening (Fig. 3B). The selected compounds were then processed by the MPNN_AAC and MPNN_Transformer models, which reduced the number of candidate chemicals to 0.0356% of the initial compounds. We also calculated interaction scores by LIGHTHOUSE and visualized them in a 2D plot (Fig. 3C, left). The best candidates would be expected to have high confidence and interaction scores, appearing in the upper right corner of the plot. Indeed, this criterion was met by several well-known drug-target combinations (Fig. 3C, right).

**Figure 3.**
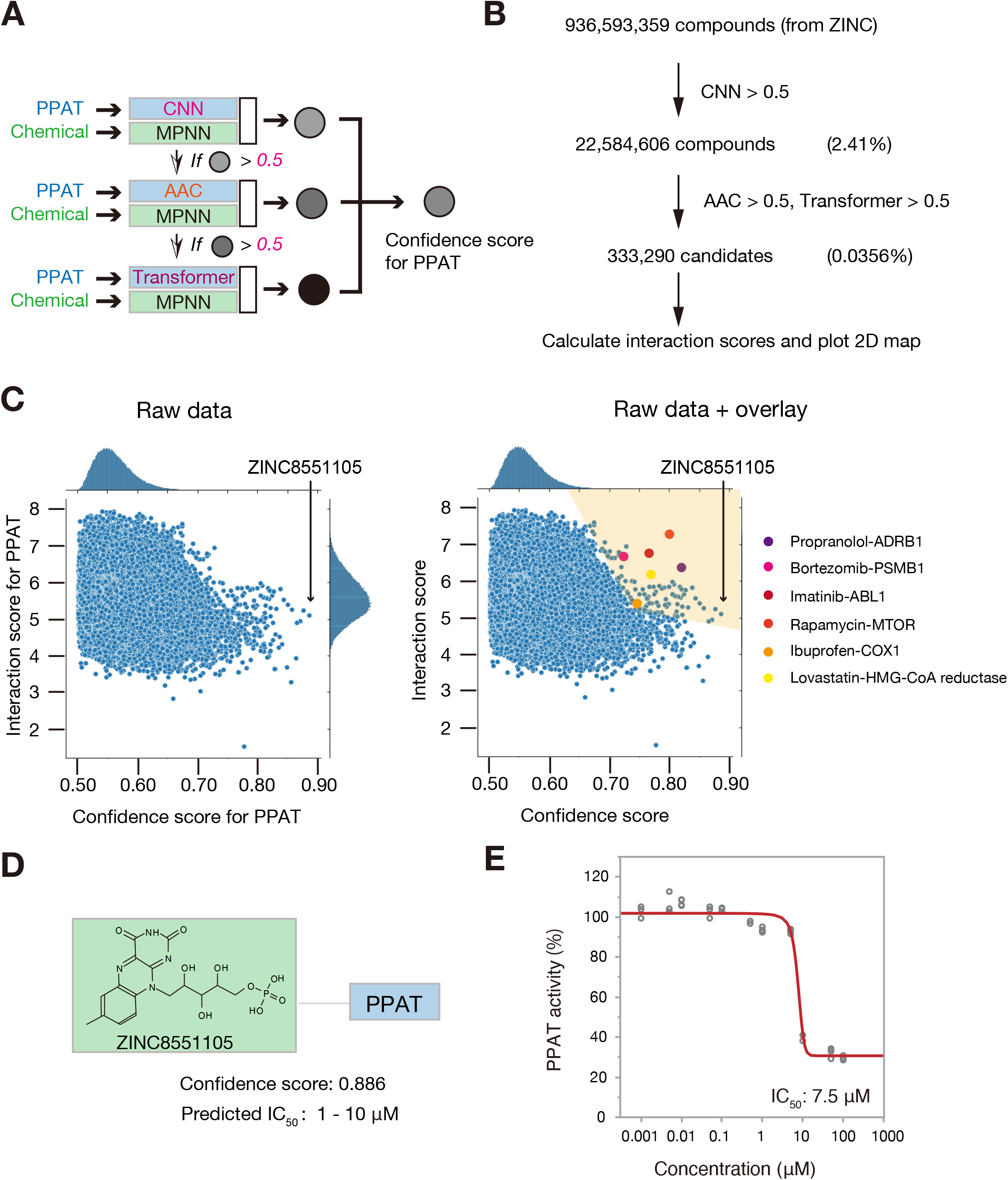
Discovery of lead compounds for treatment of cancer. (A) Scheme for PPAT inhibitor discovery. The amino acid sequence of PPAT (517 residues) and the SMILE representation for each chemical were entered into the MPNN_CNN model. If the predicted score was >0.5, the compound was entered into MPNN_AAC, and if the new predicted score was >0.5, the compound was entered into MPNN_Transformer. The harmonic mean of the three scores was then computed to obtain the confidence score. (B) Almost 1 billion compounds in the ZINC database were processed as in (A). The first filter (MPNN_CNN score > 0.5) and subsequent two filters (MPNN_AAC score > 0.5, MPNN_Transformer score > 0.5) greatly reduced the initial chemical space (to 0.0356%). The interaction scores for these selected candidates were then also calculated. (C) A 2D map of the 333,290 selected candidates from (B) is shown on the left. Ideal candidates would be expected to have high confidence and interaction scores and would be plotted in the upper right corner of the map. Indeed, some well-known drug-target pairs meet this criterion, as shown on the right, with compounds represented by the blue circles in the shaded area potentially possessing inhibitory activity for PPAT. ABL1, ABL proto-oncogene 1; COX1, cyclooxygenase 1; HMG-CoA, 3-hydroxy-3-methylyglutaryl–coenzyme A. (D) The top hit compound, ZINC8551105 (riboflavin 5’ -monophosphate), is shown together with its confidence score and estimated IC_50_ value. (E) *In vitro* PPAT activity assay performed in the presence of various concentrations (1 nM, 5 nM, 10 nM, 50 nM, 100 nM, 500 nM, 1 μM, 5 μM, 10 μM, 50 μM, and 100 μM) of riboflavin 5’ -monophosphate, with the determined IC_50_ value being within the range predicted by LIGHTHOUSE. Data are shown for four biological replicates. See also Supplementary Fig. 5.

Among the >333,000 final compounds, the top candidate PPAT inhibitor with the highest confidence score was ZINC8551105 (riboflavin 5’ -monophosphate), with a predicted IC_50_ of 1 to 10 μM (Fig. 3D). We performed a biochemical assay to test this prediction and found that riboflavin 5’ -monophosphate indeed markedly inhibited PPAT activity with an actual IC_50_ of 7.5 μM (Fig. 3E). This compound, discovered by LIGHTHOUSE solely on the basis of the PPAT amino acid sequence, is thus a potential lead compound for the development of new therapeutics targeted to a variety of cancers. It is also of note that we tested only this compound, so other top candidates might also inhibit PPAT activity.

### LIGHTHOUSE identifies an inhibitor of drug-resistant bacterial growth

Bacterial infections pose a clinical problem worldwide, especially in developing countries, and the emergence of drug-resistant bacterial strains as a result of the overuse of antibiotics has exacerbated this problem. β-Lactamase enzymes produced by antibiotic-resistant bacteria^29^ target the β-lactam ring of antibiotics of the penicillin family. We therefore applied LIGHTHOUSE to search for antibiotics not dependent on β-lactam structure.

LIGHTHOUSE predicted that pyridoxal 5’-phosphate might associate with penicillin binding proteins such as PBP2 (*mrdA*), PBP3 (*ftsI*), and PBP5 (*dacA*), all of which are essential for cell wall synthesis in *Escherichia coli*^30^ (Fig. 4A). This compound indeed suppressed the growth of wild-type *E. coli* strain JM109 in a concentration-dependent manner (Fig. 4B). Importantly, pyridoxal 5’-phosphate also markedly inhibited the growth of an ampicillin-resistant *E. coli* transformant that produces β-lactamase (Fig. 4C). These results thus suggested that, even though it was trained with human proteins, LIGHTHOUSE can also be applied to nonhuman (even bacterial) proteins.

**Figure 4.**
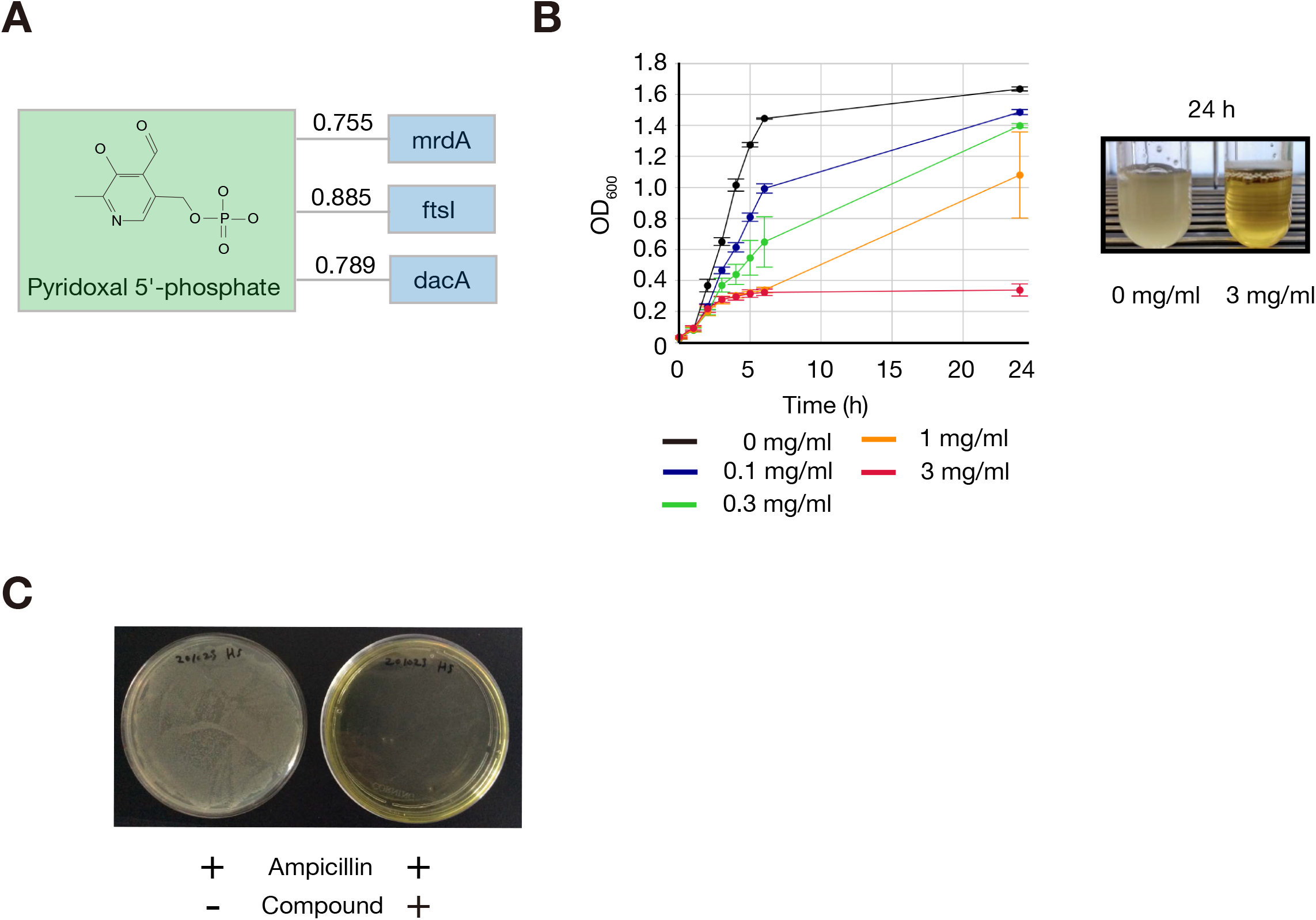
LIGHTHOUSE is applicable to prokaryotic proteins. (A) LIGHTHOUSE predicted that pyridoxal 5’ -phosphate would associate with several penicillin binding proteins (PBPs) of *E. coli* (strain K12). MrdA and FtsL are peptidoglycan D,D-transpeptidases, whereas DacA is a D-alanyl-D-alanine carboxypeptidase. (B) *Escherichia coli* strain JM109 was cultured in 2xYT medium supplemented with various concentrations of pyridoxal 5’ -phosphate, and optical density at 600 nm (OD_600_) of the culture was monitored. Data are means ± SD for three independent experiments. (C) The JM109 strain of *E. coli* was transformed with the pBlueScript II SK+ plasmid, which contains an ampicillin resistance gene as a selection marker, and the cells were plated on LB agar plates containing ampicillin in the absence or presence of pyridoxal 5’ -phosphate (3 mg/ml) and were incubated overnight. The pH of pyridoxal 5’ -phosphate was adjusted to 7.0 in order to avoid potential nonspecific toxicity.

### LIGHTHOUSE informs optimization of lead compounds

Diabetes mellitus is also a serious public health concern, with the number of affected individuals expected to increase markedly in the coming decades.^31^ Dipeptidyl peptidase–4 (DPP-4) cleaves and inactivates the incretin hormones GLP-1 and glucose-dependent insulinotropic polypeptide (GIP), and DPP-4 inhibitors are a new class of antidiabetes drug.^32^ Given that LIGHTHOUSE also predicts interaction scores, we examined whether it might also contribute to the optimization step of drug development. Indeed, LIGHTHOUSE accurately predicted the rank order of potency for several recently identified DPP-4 inhibitor derivatives^33^ (Fig. 5A). Furthermore, LIGHTHOUSE predicted that removal of the phosphate group would reduce the inhibitory potency of riboflavin 5’ -monophosphate for PPAT (Fig. 3E), and this prediction was confirmed correct by the finding that the IC_50_ value was increased from 7.5 to 49.9 µM (Fig. 5B). These data suggested the possibility that LIGHTHOUSE is capable of predicting activity cliffs.

**Figure 5.**
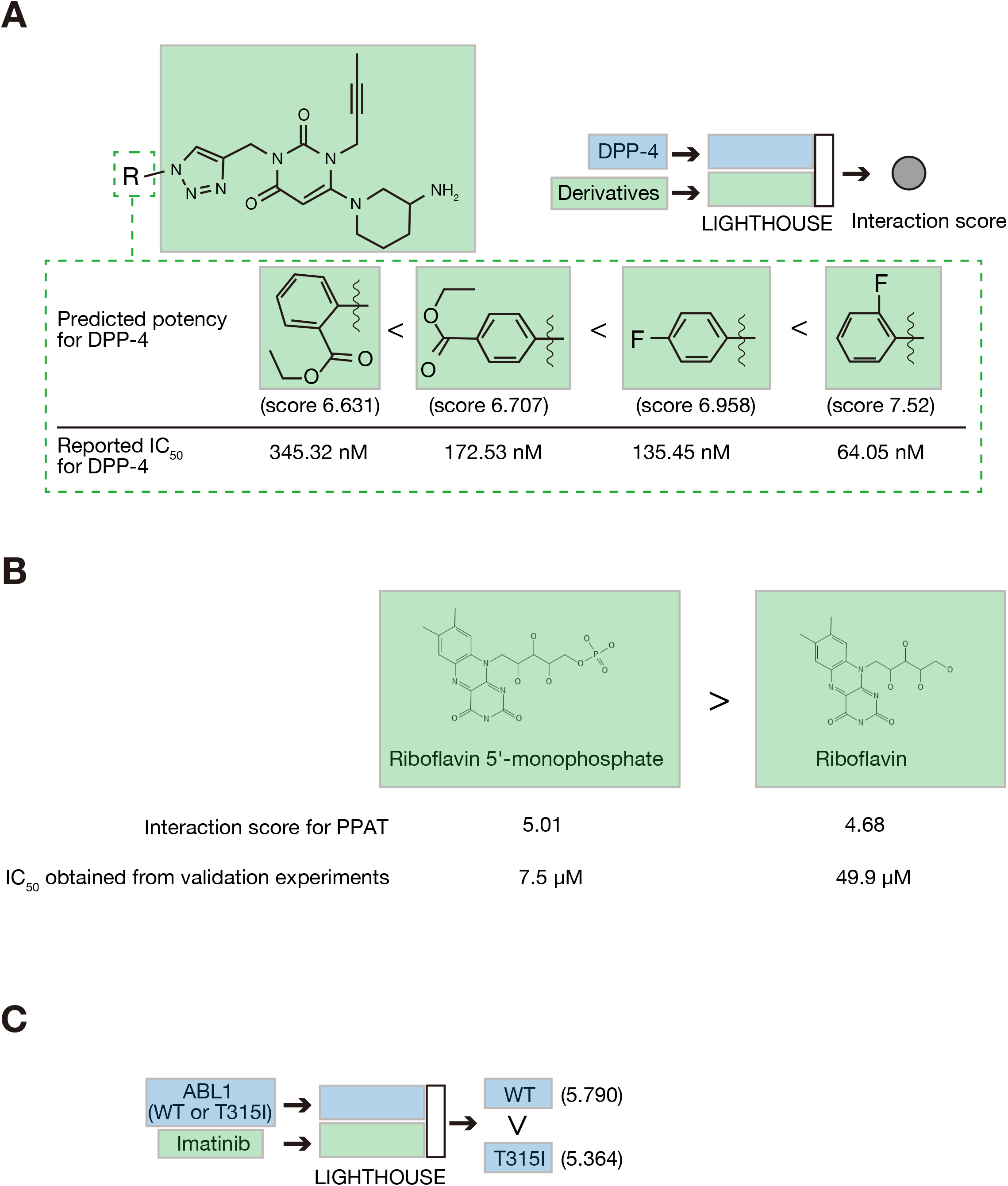
LIGHTHOUSE directs optimization of lead compounds. (A) Prediction of the potency of DPP-4 inhibitor derivatives by LIGHTHOUSE. The predicted interaction scores are compared with the reported IC_50_ values. (B) LIGHTHOUSE accurately predicts that riboflavin is a less potent PPAT inhibitor than is riboflavin 5’ -monophosphate. (C) The interaction scores calculated by LIGHTHOUSE for both wild-type (WT) and T315I mutant forms of ABL1 accurately predict that the point mutation reduces the effectiveness of the leukemia drug imatinib.

LIGHTHOUSE is also able to estimate the effect of point mutations on CPIs. For example, the T315I mutation of ABL1 in leukemia cells reduces the efficacy of imatinib,^34^ and LIGHTHOUSE accurately predicted the effect of this mutation (Fig. 5C). LIGHTHOUSE is able to provide such insight from only wild-type amino acid sequences, given the lack of variant information in the original training data set. Our results suggest that LIGHTHOUSE is able to predict the effects of small changes in protein or chemical structure, and that this will be the case even if such variants do not exist in nature.

### LIGHTHOUSE identifies potential on- and off-targets of given compounds

Opposite to the mode of drug discovery for a given protein, LIGHTHOUSE might also be able to identify proteins as potential on- or off-targets for a given compound. To verify this notion, we examined statins, which are HMG-CoA reductase inhibitors widely administered for the treatment of hyperlipidemia. Epidemiological studies have shown that statins not only lower cholesterol, however, but also have effects on cancer, although the target molecules for these effects have remained unclear.^35^ We therefore applied LIGHTHOUSE to three representative statins (atorvastatin, cerivastatin, and fluvastatin) and computed confidence scores for all human protein-coding genes (Fig. 6A and 6B, Supplementary Table 5). We then sorted the genes on the basis of these confidence scores and performed Kyoto Encyclopedia of Genes and Genomes (KEGG) pathway enrichment analysis for the top 500 potential statin targets. In addition to lipid-related pathways such as atherosclerosis and fatty liver, “pathways in cancer” was one of the most enriched KEGG pathways (Fig. 6C), consistent with previous findings.^36–39^ Potential targets of statins for cancer treatment identified by LIGHTHOUSE included STAT3, CCND1, AKT1, and CCL2 (Fig. 6D).

**Figure 6.**
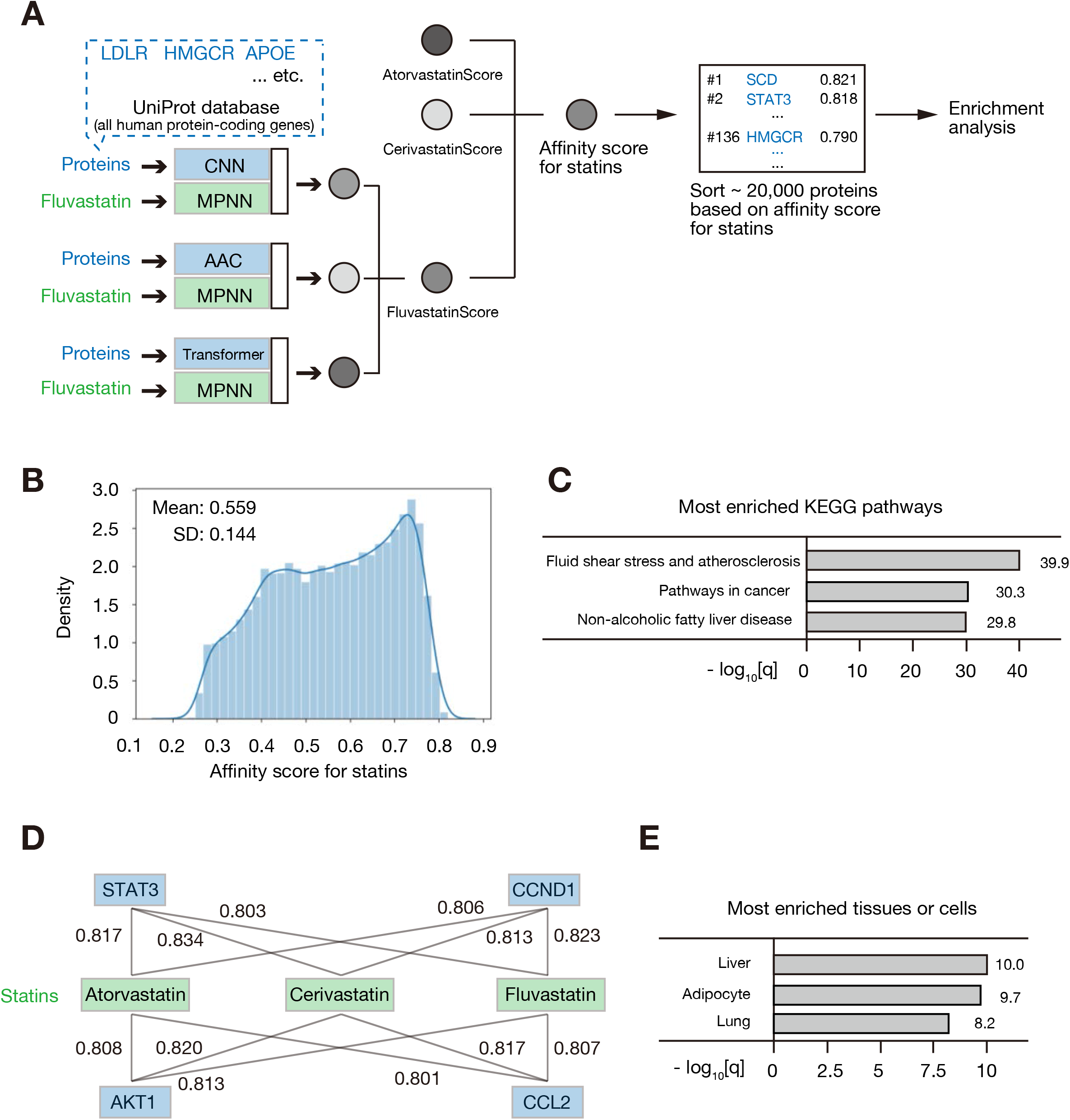
LIGHTHOUSE uncovers potential target proteins for given drugs. (A) Identification of statin targets by LIGHTHOUSE. LIGHTHOUSE was applied to calculate confidence scores for all human protein-coding genes in the UniProt database and fluvastatin, atorvastatin, and cerivastatin. The harmonic mean of these confidence scores (FluvastatinScore, AtorvastatinScore, and CerivastatinScore) was calculated as an affinity score for statins. Sorting on the basis of this affinity score yielded a list of potential statin target proteins. HMGCR (HMG-CoA reductase), a known key target of statins, was ranked 136th with a score of 0.790. The top 500 identified genes were then subjected to enrichment analysis. LDLR, low-density lipoprotein receptor; APOE, apolipoprotein E; SCD, stearoyl-CoA desaturase; STAT3, signal transducer and activator of transcription 3. (B) Distribution of the harmonic mean of the AtorvastatinScore, CerivastatinScore, and FluvastatinScore (affinity score for statins). (C) KEGG pathway enrichment analysis for the top 500 potential statin targets identified by LIGHTHOUSE. Minus log_10_-transformed q values are shown. (D) Confidence scores for representative predictions by LIGHTHOUSE of the association of statins with cancer-related proteins. (E) Enrichment analysis for expression sites of the top 500 potential statin targets. Minus log_10_-transformed q values are shown. See also Supplementary Table 5.

Given that side effects of drugs often manifest in organs that express target proteins, we hypothesized that LIGHTHOUSE might be able to identify which organs are at risk of damage from a given drug. We performed another enrichment analysis for the same top 500 potential statin target genes to determine which organs or cell types preferentially express these genes. The top three candidates were the liver, adipocytes, and lung (Fig. 6E), consistent with the liver being the primary site of statin metabolism and interstitial pneumonia being one of the most severe side effects of statins.^40^ Prediction of potential target proteins for a given drug by LIGHTHOUSE will thus provide insight into which organs warrant close monitoring by physicians during treatment with the drug, especially in first-in-human clinical trials.

### LIGHTHOUSE identifies novel potential therapeutics for COVID-19

SARS-CoV-2 emerged at the end of 2019 and has caused a pandemic of infectious pulmonary disease, COVID-19.^41^ We noticed that genes whose expression is up-regulated after SARS-CoV-2 infection^42–44^ were enriched in the list of potential statin targets identified by LIGHTHOUSE (Fig. 7A). Indeed, previous studies have shown that statins prevent exacerbation of COVID-19.^45,46^ With this finding that LIGHTHOUSE is also effective for COVID-19 drug discovery, we applied it to the virtual screening of ∼10,000 approved drugs, given that drug repurposing may allow faster delivery of effective agents to patients in need. We calculated scores for angiotensin-converting enzyme 2 (ACE2), which is targeted by SARS-CoV-2 for infection of host cells,^47^ and the top drug candidate, ethoxzolamide, was selected for validation analysis (Fig. 7B). Immunocytofluorescence analysis revealed that ethoxzolamide blocks proliferation of SARS-CoV-2 in Vero-TMPRSS2 cells (Fig. 7C). Furthermore, ethoxzolamide was effective against not only the wild-type virus but also the alpha, beta, gamma, and delta variants. It thus rescued virus-challenged cells in a concentration-dependent manner without affecting noninfected cells (median cytotoxicity concentration > 50 μM) (Fig. 7D and 7E, Supplementary Fig. 6, Supplementary Table 6), and it reduced the virus load present in the culture supernatant of the cells (Fig. 7F and 7G, Supplementary Fig. 7). Ethoxzolamide is approved for the treatment of seizures and glaucoma,^48,49^ and its pharmacodynamics are therefore known. It is therefore immediately available for repurposing for the treatment of patients with COVID-19, with its further optimization having the potential to save many lives. We tested another 11 drugs and found another 2 compounds that also inhibited SARS-CoV-2 infection (Supplementary Table 7). This high hit rate of 25% (3 hits in 12 compounds) shows the potential power of LIGHTHOUSE for drug repositioning.

**Figure 7.**
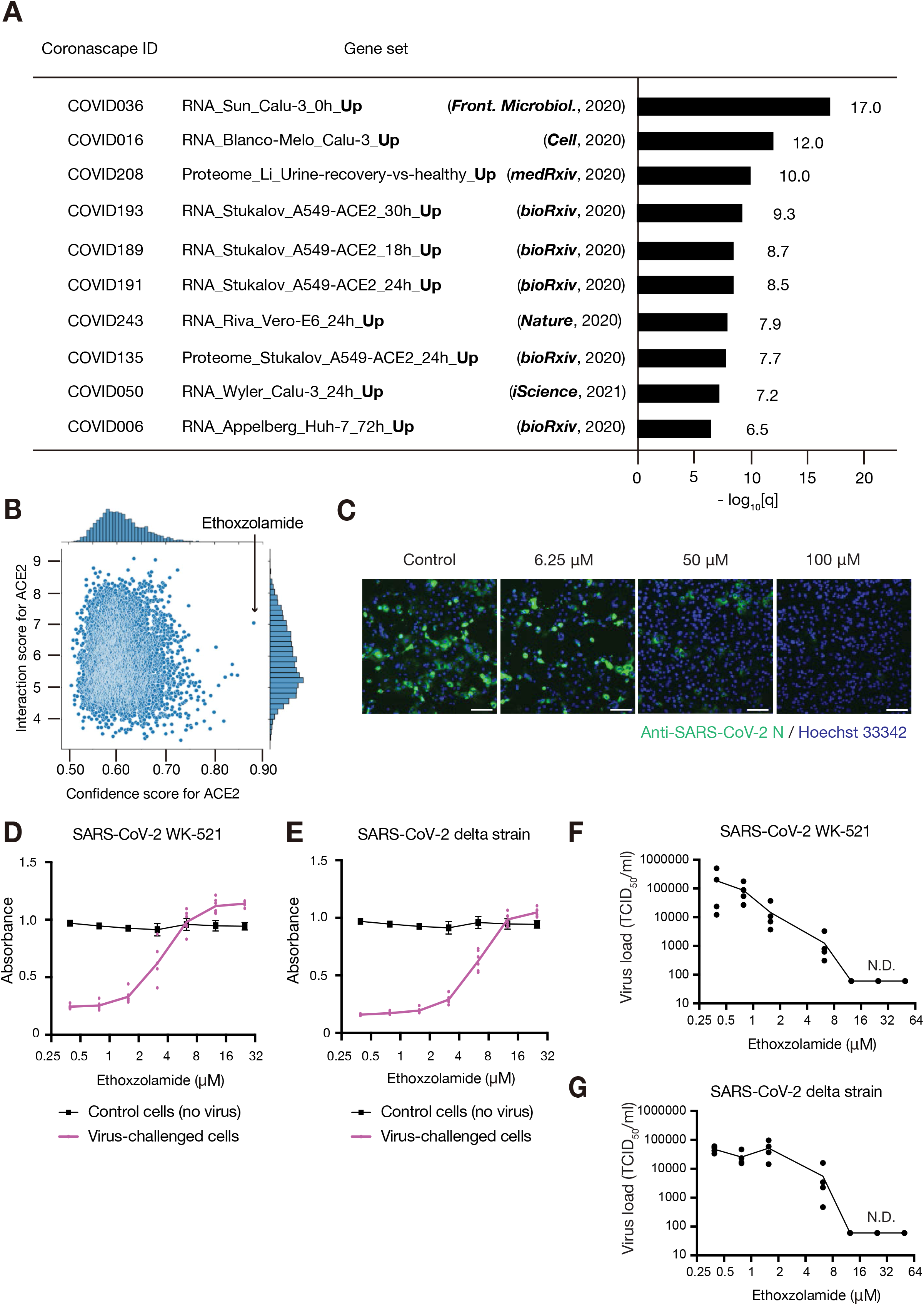
LIGHTHOUSE-based drug repurposing for COVID-19. (A) Enrichment analysis of the top 500 potential statin targets identified in Fig. 6 for COVID-19–associated gene sets. Minus log_10_-transformed q values are shown. (B) Prediction by LIGHTHOUSE of ethoxzolamide as a potential therapeutic for SARS-CoV-2 infection on the basis of its confidence and interaction scores for ACE2. Vero-TMPRSS2 cells were infected with wild-type SARS-CoV-2 at an MOI of 0.0001, (C) cultured for 64 h in the presence of the indicated concentrations of ethoxzolamide, and subjected to immunocytofluorescence analysis with antibodies to SARS-CoV-2 N protein (green). Nuclei were stained with Hoechst 33342 (blue). Scale bars, 100 µm. (D and E) Vero-TMPRSS2 cells challenged with wild-type (WK-521) or delta strains of SARS-CoV-2, respectively, were cultured in the presence of various concentrations of ethoxzolamide for 3 days and then subjected to the MTT assay of cell viability. Nonchallenged cells were examined as a control. Data are means ± SD for three independent experiments each performed in duplicate. (F and G) Effect of ethoxzolamide on the SARS-CoV-2 load in culture supernatants of Vero-TMPRSS2 cells challenged with wild-type (WK-521) or delta strains of the virus, respectively. Data are from four independent experiments, with the graph line connecting mean values. TCID_50_, median tissue culture infectious dose; N.D., not detected. See also Supplementary Figs. 6, 7, Supplementary Table 6.

## Discussion

Although recent advances in biological and medical research have uncovered various proteins as promising therapeutic targets in a variety of diseases, the clinical application of these research findings has been limited because of the difficulty in identifying therapeutic chemicals for these targets in a cost-effective and high-throughput manner. Acquisition of 3D structural data for target proteins has been labor-intensive, and processing of such data requires a huge amount of computer capacity and time, resulting in a delay in the translation of research findings from the laboratory to the clinic. We have now shown that LIGHTHOUSE facilitates the identification, from a vast chemical space, of candidate compounds for given target proteins solely on the basis of the primary structure of these proteins. Furthermore, the AUROC for LIGHTHOUSE is equivalent to or better than that for state-of-the-art 3D docking simulation methods as well as for other AI methodologies.

We have applied LIGHTHOUSE to attractive targets for various diseases, including cancer, bacterial infection, metabolic diseases, and COVID-19, with some of the suggested chemicals being determined experimentally to be effective for inhibition of the corresponding targets. The actual targets of identified compounds require further biological investigation, especially in the case of pyridoxal 5’-phosphate and PBPs. As for ethoxzolamide, we attempted to obtain clinical evidence in support of its effectiveness against COVID-19. However, ethoxzolamide is an old drug and is essentially no longer prescribed as a result of the development of more potent agents such as acetazolamide. We were therefore not able to find data to address whether COVID-19 patients taking ethoxzolamide have a better clinical outcome.

The hit compounds themselves identified in the present study are not sufficient for immediate clinical use. LIGHTHOUSE can be applied not only for the identification of lead compounds but also for their subsequent optimization, which requires extensive work to identify more potent and specific or less toxic derivatives. One promising method to support such optimization is to apply LIGHTHOUSE and either reinforcement learning^50^ (Supplementary Fig. 8) or Metropolis-Hasting (MH) approaches together. Virtual libraries can be generated from identified lead compounds in an intensive manner with the use of sophisticated chemoinformatics algorithms such as RECAP (Retrosynthetic Combinatorial Analysis Procedure).^51^ LIGHTHOUSE can score the generated virtual compounds and help to narrow down the candidates with better scores than the original hit compound. Select candidates can then be synthesized in collaboration with organic chemists and their effects tested. Given the recent success of the MH approach in various life science fields,^52,53^ LIGHTHOUSE should be able to facilitate the optimization of drug candidates by iterating these steps.

The biggest limitation of LIGHTHOUSE is the generation of false positives, which is due in part to the fact that the confidence score provided by STITCH is not based solely on experimental data but also on other factors such as co-occurrence in the literature. This confidence score therefore does not necessarily reflect actual interaction for a given protein-chemical pair, and well-studied molecules are thus prone to score higher than others. On the other hand, low confidence scores do not necessarily mean that the protein and chemical in question do not interact. We modeled this score because current biologically determined CPI data sets contain fewer CPI pairs. In addition, as a result of publication bias and biological experimental conditions, it is sometimes difficult to tell whether a protein and chemical do not interact, whether they did not bind under the specific assay condition, or whether they were not tested. This drawback of LIGHTHOUSE can be mitigated partially by combining the three different models (CNN, AAC, and Transformer). It may also be important to perform a counter–virtual screening to determine whether an identified small molecule reacts specifically with the target protein or whether it scores highly with many proteins. Such an approach has the potential to reduce the number of false positives and provide more accurate guidance. Furthermore, we also modeled IC_50_ data from BindingDB, which is derived from actual bioassays. We found that the predicted value and observed value correlate well in the BindingDB test data set (Supplementary Fig. 4). By combining the confidence score (from STITCH) and the interaction score (from BindingDB) provided by LIGHTHOUSE, we were able to discover a clinically approved drug that was able to block SARS-CoV-2 infection.

Another potential limitation is that the performance for STITCH data might be overestimated because we downsampled the original imbalanced data (Supplementary Table 1). There are several ways to tackle imbalanced data, including downsampling, oversampling, and more complex sampling methods such as SMOTE.^54^ Each of these approaches has potential limitations, however.^55,56^ In the present study, we adopted downsampling, as in a previous study,^7^ so the performance for STITCH data may be overestimated when compared with use of the entire STITCH data.

Despite these limitations, LIGHTHOUSE proved to be effective for the identification of lead compounds for all conditions tested. It can theoretically be applied to any protein of any organism, and even to proteins that do not exist naturally. This is an advantage over 3D docking simulation methods, which require prior 3D structural knowledge of the protein of interest. LIGHTHOUSE computes and embeds structural information in numerical vectors, which are then readily retrieved by the subsequent decoding module. Given the accelerating development of protein embedding technologies^57^ and graph-based chemoinformatics approaches, LIGHTHOUSE has the potential to be a cornerstone of drug discovery. It is also of note that we split the STITCH data set once, given that a previous study^7^ showed it could obtain high performance with a relatively small range of hyperparameter tuning. In-depth hyperparameter tuning with cross-validation may further boost the performance of the model.

The next step for more sophisticated AI-dependent drug discovery and its clinical application will be the use of huge-scale and robust protein-chemical binding (and nonbinding) data, with the recent introduction of academic journals that specialize in big data reflecting the growth of this field. Moreover, development of improved approaches to the handling of imbalanced data^58^ is another active research field in computer science. These advances in both data and methodology should allow more reliable modeling of physical interactions and further facilitate drug discovery in the coming years.

In summary, we have developed LIGHTHOUSE as a means to discover promising lead compounds for any target protein irrespective of its 3D structural information. Furthermore, we have demonstrated the power of LIGHTHOUSE by identifying and validating novel therapeutics for various global health concerns including COVID-19. LIGHTHOUSE will serve as a guide for researchers in all areas of biomedicine, paving the way for a wide range of future applications.

## EXPERIMENTAL PROCEDURES

### Resource availability

#### Lead contact

Further information and requests for resources and reagents should be directed to and will be fulfilled by the lead contact, Keiichi I. Nakayama.

#### Materials availability

This study did not generate new unique reagents.

#### Data and code availability

All data used for training are downaloaded and publicity available from STITCH (http://stitch.embl.de)^14^ and BindingDB (https://www.bindingdb.org/bind/index.jsp)^18^ web servers. SMILES representations for small-molecule compounds were downloaded from PubChem (https://pubchem.ncbi.nlm.nih.gov) or ZINC15 (https://zinc15.docking.org).^28^ Amino acid sequences were obtained from UniProt (https://www.uniprot.org).^59^ For drug repurposing analyses, we used the KEGG-DRUG database (https://www.genome.jp/kegg/drug).^60^ All referenced COVID-19 signatures are available at Coronascape (https://metascape.org/COVID).^61^

Code for LIGHTHOUSE with pretrained weights together with a notebook reproducing the results presented in this paper is available at https://github.com/Shimizu-Lab/LIGHTHOUSE.

### Generation of a data set for the training phase of LIGHTHOUSE

The compound SMILES strings of the data set were extracted from the PubChem compound database on the basis of compound names and PubChem compound IDs (CIDs). The protein sequences of the data set were extracted from the UniProt protein database on the basis of gene names/RefSeq accession numbers or the UniProt IDs. We downloaded the protein-chemical link data set of *Homo sapiens* (Taxonomy ID 9606) from the STITCH database (version 5.0). Given that the STITCH score is heavily biased toward 0, we separated the data into nine bins on the basis of the score and stratify-extracted the same number of CPIs (140,000 each), yielding 1,260,000 CPIs (Supplementary Table 1). We then randomly separated these data into training (80%), validation (10%), and test (10%) data sets (Supplementary Fig. 1A). With regard to IC_50_, we downloaded data from BindingDB, obtained SMILES expressions and amino acid sequences similarly, and again separated the data into training (80%), validation (10%), and test (10%) data sets (Supplementary Fig. 2A). Given that IC_50_ values differ widely, we scaled the values by log transformation (Eq. 1) and used the transformed values for BindingDB training.

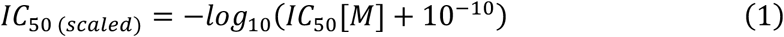

### LIGHTHOUSE architecture and training

The proposed overall model comprises two encoder networks (for chemicals and proteins) and one decoder network. MPNN is a message passing graph neural network that operates on compound molecular graphs.^11^ In brief, MPNN conveys latent information among the atoms and edges. The message passing phase runs for *t* time steps and is defined in terms of message functions *M*_*t*_ and vertex update functions *U*_*t*_. During this phase, hidden states 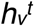 (128 dimensions in our model) at each node in the chemical graph are updated with the incoming messages 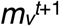 according to the following equations (Eqs. 2 and 3):

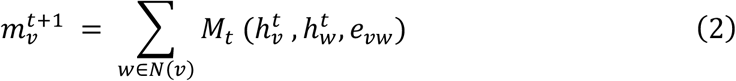

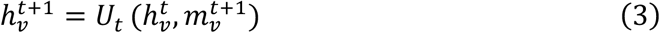

where *e*_*vw*_ represents edge feature between nodes *v* and *w, N*(*v*) denotes the neighbor nodes of vertex *v* in graph *G*, and message functions *M*_*t*_ and update functions *U*_*t*_ are learned differentiable functions. After *T* (= 3) cycles of message passing and subsequent update, a readout function (average) is used to extract the embedding vectors at the graph level.

CNN is powerful for computer vision, but here we used a multilayer 1D CNN for protein sequence, as described previously.^6^ In brief, the target amino acid is decomposed to each individual character and is encoded with an embedding layer and then fed into the CNN convolutions. We used three consecutive 1D convolutional layers with an increasing number of filters, with the second layer having double and the third layer having triple the number of filters in the first layer (32, 64, and 96 filters for the three layers). The convolution layers are followed by a global max-pooling layer. AAC is an 8420-length vector in which each position corresponds to a sequence of three amino acids.^13^ Transformer uses a self-attention–based transformer encoder^12^ that operates on the substructure partition fingerprint of proteins. Algorithmically speaking, Transformer follows *O*(*n*^4^) in computation time and memory, where *n* is the input size.^12^ This bottleneck prevented us from considering each amino acid as a token. We therefore used partition fingerprints to decompose amino acid sequence into protein substructures of moderate size and then fed each of the partitions into the model as a token.^8^

As for the decoder, we exploited a previously described architecture.^4^ In brief, encoder outputs are concatenated and entered into a three-layer feed-forward dense neural network (1024,1024, and 512 nodes), which finally outputs one value. We used Rectified Linear Unit (ReLU),^62^ *g*(*x*) = max(0,*x*), as the activation function in the decoder network.

We defined our loss function with MSE (Eq. 4):

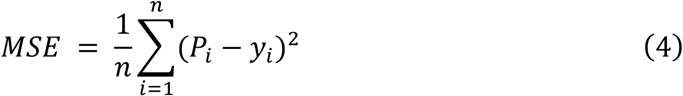

where *P*_*i*_ is the LIGHTHOUSE-predicted score for the *i*th compound-protein pair and *Y*_*i*_ is the true label in the corresponding training data, with a batch size of 128. We trained three architectures (MPNN_CNN, MPNN_AAC, MPNN_Transformer) separately for the STITCH and BindingDB training data with the Adam optimizer and a learning rate of 0.001. For evaluation metrics, we used MSE, concordance index, and Pearson correlation as well as AUROC. For every 10 epochs, we compared the current loss (in the validation data set) with that of 10 epochs ago; if the loss was not decreasing, we terminated the training for that model. As a result of this early termination, we trained MPNN_CNN for 40 epochs, MPNN_AAC for 70 epochs, and MPNN_Transformer for 100 epochs with regard to the confidence score (Supplementary Fig. 1B–1G). As for the models for the interaction score, we trained MPNN_CNN for 70 epochs, MPNN_AAC for 100 epochs, and MPNN_Transformer for 70 epochs (Supplementary Fig. 2B–2G), according to the same guidelines. After the training was completed, we finally evaluated the models with the test data sets, which were kept aside during the training and so had not previously been seen by the models.

### Generation of virtual chemical libraries and prediction by LIGHTHOUSE

We prepared nearly 1 billion purchasable substances, which were downloaded from the ZINC database^28^ as of 30 July 2020, for virtual PPAT inhibitor screening. For drug repurposing, we obtained approved drugs from the KEGG-DRUG database^60^ as of 24 January 2021. For calculation of confidence and interaction scores, we fixed the proteins of interest (PPAT or ACE2) and changed the compounds iteratively, which yielded lists of predicted scores for all the compounds tested. As for the peptide drugs shown in Fig. 2, we converted them as for small-molecule compounds with the use of SMILES.

### PPAT activity assay

Sf21 cells were cultured in Sf-900 II SFM (Gibco, Cat# 10902-088) supplemented with 10 μM ferric ammonium citrate. They were transfected with a bacmid encoding human PPAT for 64 h, harvested, washed three times with phosphate-buffered saline, and lysed in a solution containing 150 mM NaCl, 25 mM Tris-HCl (pH 7.4), 0.5% Triton X-100, and 5 mM EDTA. The lysate was centrifuged at 10,000 × *g* for 6 min at 4°C, and the resulting supernatant (100 ng/ml) was incubated for 4 h at 37°C together with 5 mM glutamine (Gibco, Cat# 25030-081), 1 mM phosphoribosyl pyrophosphate (Sigma, Cat# P8296), 10 mM MgCl_2_, 50 mM Tris-HCl (pH 7.4), and various concentrations of riboflavin 5’ -monophosphate (Sigma, Cat# F2253-10 mg). Enzyme activity was assessed on the basis of glutamate production as measured with a glutamate assay kit (Abcam, Cat# 138883). The IC_50_ value was estimated from biological quadruplicates with a four-parameter logistic model^63^ and with the use of JMP pro 15 software (version 15.1.0).

### Assay of *E. coli* growth

Portions (20 µl) of *E. coli* strain JM109 (1 × 10^10^ colony-forming units (CFU)/ml) were cultured for various times in 2 ml of 2xYT liquid medium (BD Difco, Cat# 244020) containing various concentrations of pyridoxal 5’ -phosphate (pH adjusted to 7.0), after which OD_600_ was measured with a GENESYS 30 visible spectrophotometer (ThermoFisher Scientific, Cat# 840-277000). In addition, the JM109 strain was transformed with 1 μg of the pBlueScript II SK+ plasmid (Invitrogen), which harbors an ampicillin resistance gene as a selection marker, and was then spread on LB agar plates containing ampicillin (100 μg/ml) (Wako, Cat# 012-23303) with or without pyridoxal 5’ -phosphate (3 mg/ml) and incubated overnight.

### Virtual identification of statin targets and enrichment analyses

Three representative statins were fixed as chemical inputs, and all human protein-coding genes in the UniProt database were iteratively changed. The harmonic mean of the three confidence scores was calculated as an affinity score for statins, and the human protein-coding genes were sorted on the basis of this score. The resulting top 500 potential targets were then subjected to enrichment analyses with the use of the Metascape web server.^61^

### SARS-CoV-2 assays

Vero-TMPRSS2 cells^64^ were maintained in Dulbecco’s modified Eagle’s medium (DMEM) supplemented with 10% fetal bovine serum. The WK-521 strain of SARS-CoV-2 (EPI_ISL_408667) as well as the alpha (QK002, EPI_ISL_768526), beta (TY7-501, EPI_ISL_833366), gamma (TY8-612, EPI_ISL_1123289), and delta (TY11-927, EPI_ISL_2158617) variants were obtained from National Institute of Infectious Diseases in Japan. Stocks of these viruses were prepared by inoculation of Vero-TMPRSS2 cell cultures as described previously.^64^ The MTT assay was performed to evaluate cell viability after virus infection also as previously described.^64^ In brief, serial twofold dilutions of ethoxzolamide in minimum essential medium (MEM) supplemented with 2% fetal bovine serum were added in duplicate to 96-well microplates. Vero-TMPRSS2 cells infected with wild-type or variant SARS-CoV-2 at 4 to 10 TCID_50_ (median tissue culture infectious dose) were also added to the plates, which were then incubated at 37°C for 3 days. The viability of the cells was then determined with the MTT assay, and the culture supernatants were harvested for determination of the TCID_50_ value as a measure of viral load. For indirect immunofluorescence analysis, cells infected with wild-type SARS-CoV-2 at a multiplicity of infection (MOI) of 0.0001 were cultured in the presence of various concentrations of ethoxzolamide for 64 h, fixed with 3.7% buffered formaldehyde, permeabilized with 0.05% Triton X-100, and incubated with antibodies to SARS-CoV-2 N protein (GeneTex, Cat# GTX635679). Immune complexes were detected with Alexa Fluor Plus 488–conjugated goat antibodies to rabbit immunoglobulin G (Invitrogen–ThermoFisher Scientific, Cat# A32731). Nuclei were stained with Hoechst 33342 (Invitrogen). Fluorescence images were captured with an IX73 fluorescence microscope (Olympus).

## SUPPLEMENTAL INFORMATION

Supplemental information can be found onine at….

## ACKNOWLEDGMENTS

This work was supported by KAKENHI grants from the Japan Society for the Promotion of Science (JSPS) to H. Shimizu (JP21K17856), M.K. (JP21K15068), and K.I.N. (JP18H05215), as well as by a grant from the Japan Agency for Medical Research and Development (AMED) to K.I.N. (21wm0425002). H. Shimizu was also supported by postdoctoral fellowships from Takeda Science Foundation and Mochida Foundation. We thank T. Sawada and R. Wada for technical assistance, P.A. Silver and J.C. Way for critical reading of the manuscript, other laboratory members for discussion, and A. Ohta for help with preparation of the manuscript.

## AUTHOR CONTRIBUTIONS

H. Shimizu conceived of and designed the study, developed LIGHTHOUSE, performed validation experiments, and wrote the original draft of the manuscript. M.K. conducted PPAT validation assays. Y.O., M.S., A.S., and H. Sawa performed COVID-19 infection analyses. M.M. contributed to discussion. K.I.N. supervised the study and edited the manuscript.

