## Supplemental Information for "LIGHTHOUSE illuminates therapeutics for a variety of diseases including COVID-19"

**SUPPLEMENTAL TABLE TITLES**
**SUPPLEMENTAL TABLES 1–4 AND 6–7 (SUPPLEMENTAL TABLE 5 IS PROVIDED SEPARATELY IN CSV FORMAT)**
**SUPPLEMENTAL FIGURE LEGENDS**
**SUPPLEMENTAL FIGURES 1–8**

**SUPPLEMENTAL TABLE TITLES**

**Table S1.** Stratified sampling of the STITCH database.

**Table S2.** Final performance of LIGHTHOUSE for confidence scores with test data.

**Table S3.** Final performance of LIGHTHOUSE for interaction scores with test data.

**Table S4.** Comparison of model performance for LIGHTHOUSE and other state-of-the-art methods.

**Table S5.** Confidence scores for statins. (PROVIDED SEPARATELY IN CSV FORMAT)

**Table S6.** Median effective concentration ($EC_{50}$) values of ethoxzolamide for Vero-TMPRSS2 cells challenged with various SARS-CoV-2 strains.

**Table S7.** Median effective concentration ($EC_{50}$), median cytotoxic concentration ($CC_{50}$), and selectivity index of 12 approved drugs for Vero-TMPRSS2 cells challenged with various SARS-CoV-2 strains.

| Score | STITCH original | Stratified sampling |
|---|---|---|
| 0.0 ~ 0.2 | 5,310,242 | 140,000 |
| 0.2 ~ 0.3 | 5,237,589 | 140,000 |
| 0.3 ~ 0.4 | 1,769,008 | 140,000 |
| 0.4 ~ 0.5 | 507,870 | 140,000 |
| 0.5 ~ 0.6 | 264,566 | 140,000 |
| 0.6 ~ 0.7 | 200,401 | 140,000 |
| 0.7 ~ 0.8 | 161,656 | 140,000 |
| 0.8 ~ 0.9 | 143,333 | 140,000 |
| 0.9 ~ 1.0 | 169,391 | 140,000 |
| | 13,764,056 | 1,260,000 |

**Shimizu et al., Table S1**

Performances for confidence score

| | MPNN_CNN | MPNN_AAC | MPNN_Transformer |
|---|---|---|---|
| MSE (test data) | 0.0222 | 0.0207 | 0.0195 |
| AUROC (test data) | 0.8082 | 0.8212 | 0.8280 |

**Shimizu et al., Table S2**

Perfomances for interaction score

| | MPNN_CNN | MPNN_AAC | MPNN_Transformer |
|---|---|---|---|
| MSE (test data) | 0.5820 | 0.5764 | 0.5795 |
| AUROC (test data) | 0.8432 | 0.8437 | 0.8428 |

**Shimizu et al., Table S3**

Perfomances for the dataset Tsubaki et al. provided
(https://github.com/masashitsubaki/CPI_prediction/tree/master/dataset)

| | Graph-based deep learning methods | |
|---|---|---|
| | Tsubaki et al. | LIGHTHOUSE |
| **Human** | | |
| AUROC | 0.970 | **0.991** |
| F1 | 0.920 | **0.957** |
| **C. elegans** | | |
| AUROC | 0.978 | **0.989** |
| F1 | 0.933 | **0.952** |

**Shimizu et al., Table S4**

**(SUPPLEMENTARY TABLE 5 IS PROVIDED SEPARATELY AS A CSV FORMAT)**

| | EC50 (µM) of Ethoxzolamide (mean ± SD) |
|---|---|
| SARS-CoV-2 Wuhan | 3.71 ± 0.709 |
| SARS-CoV-2 U.K. | 3.14 ± 0.429 |
| SARS-CoV-2 South Africa | 14.5 ± 0.689 |
| SARS-CoV-2 Brazil | 18.5 ± 1.74 |
| SARS-CoV-2 India | 6.18 ± 0.738 |

**Shimizu et al., Table S6**

| | | VeroE6T cells | | |
|---|---|---|---|---|
| Supplier | Catalog number | EC50 (nM) | CC50 (nM) | Selectivity Index |
| **Sigma-Aldrich** | **333328-1G** | **1698** | **62192** | **37** |
| TCI | A2598 | >100000 | | |
| **Sigma-Aldrich** | **D4628-1G** | **5695** | **12609** | **2.2** |
| Sigma-Aldrich | N5535-100G | >100000 | | |
| Sigma-Aldrich | D150959-5G | >100000 | | |
| TCI | A2476 | >100000 | | |
| **TCI** | **R0064** | **9579** | **41705** | **4.4** |
| TCI | G0392 | >100000 | | |
| Sigma-Aldrich | 46959-U | >100000 | | |
| Sigma-Aldrich | T3125-25MG | >100000 | | |
| Sigma-Aldrich | C7869-1MG | >100000 | | |
| Funakoshi | CS-0153 | >100000 | | |

**Shimizu et al., Table S7**

### SUPPLEMENTAL FIGURE LEGENDS

**Figure S1. Training process for determination of the confidence score by LIGHTHOUSE**

(A) More than 1 million chemical–(human) protein interactions (CPIs) were extracted from the STITCH database (http://stitch.embl.de) and were divided into a training set (80%), a validation set (10%), and a test set (10%). The registered confidence scores ranged from 0.15 to 1 and were calculated on the basis of experimental data, evolutionary relations, co-presentation in PubMed abstracts, and other factors. After one epoch of training, the validation data were applied. After every 10 epochs, the current loss (validation data set) was compared with the loss of 10 epochs ago to determine whether additional training was necessary. Finally, the test data were applied to evaluate model performance.

(B and C) MSE (B) and AUROC (C) for the validation data set with MPNN_CNN, which uses MPNN as the chemical encoder and CNN as the protein encoder.

(D and E) MSE (D) and AUROC (E) for the validation data set with MPNN_AAC, which uses MPNN as the chemical encoder and AAC as the protein encoder.

(F and G) MSE (F) and AUROC (G) for the validation data set with MPNN_Transformer, which uses MPNN as the chemical encoder and Transformer as the protein encoder. See also Figure 1.

**Figure S2. Training process for determination of the interaction score by LIGHTHOUSE**

(A) More than 1 million reported $IC_{50}$ values were extracted from the BindingDB database (https://www.bindingdb.org/bind/index.jsp) and were split into training (80%), validation (10%), and test (10%) data sets. Given that these values range widely, they were log-transformed. After one epoch of training, the validation data were applied. After every 10 epochs, the current loss (validation data set) was compared with the loss of 10 epochs ago to determine whether further training was necessary. The test data were finally examined to evaluate model performance.

(B and C) MSE (B) and AUROC (C) for the validation data set with MPNN_CNN, which uses MPNN as the chemical encoder and CNN as the protein encoder.

(D and E) MSE (D) and AUROC (E) for the validation data set with MPNN_AAC, which uses MPNN as the chemical encoder and AAC as the protein encoder.

(F and G) MSE (F) and AUROC (G) for the validation data set with MPNN_Transformer, which uses MPNN as the chemical encoder and Transformer as the protein encoder. See also Figure 1.

**Figure S3. LIGHTHOUSE outperforms other state-of-the-art methods with**

**DUD-E data**

(A) DUD-E data were randomly split into training (72 proteins) and test (30 proteins) data. The LIGHTHOUSE architecture was trained with the DUD-E training data set, and its performance was then evaluated with the test data set.
(B) ROC curve for LIGHTHOUSE with the DUD-E test data.
(C) Comparison of the performance of LIGHTHOUSE with that of other state-of-the-art methods including graph-based deep learning model (Tsubaki et al.). Each model was trained with the same DUD-E training data, and the AUROC values for the DUD-E test data were compared.

**Figure S4. LIGHTHOUSE also precisely predicts IC50 values**

Interaction score predicted by LIGHTHOUSE and actual $IC_{50}$ values are highly correlated (r = 0.886) for BindingDB test data.

**Figure S5. Prediction of potential PPAT inhibitors**

The amino acid sequence of PPAT and the SMILE representation of each compound in the ZINC database were entered into LIGHTHOUSE for virtual screening. See also Figure 3.

**Figure S6. Ethoxzolamide rescues cells infected with SARS-CoV-2**

Vero-TMPRSS2 cells challenged with alpha (A), beta (B), or gamma (C) strains of SARS-CoV-2 were cultured in the presence of various concentrations of ethoxzolamide for 3 days and then subjected to the MTT assay of cell viability. Nonchallenged cells were examined as a control. Data are means ± SD for three independent experiments. See also Figure 7.

**Figure S7. Ethoxzolamide reduces SARS-CoV-2 virus load**

Vero-TMPRSS2 cells challenged with alpha (A), beta (B), or gamma (C) strains of SARS-CoV-2 were cultured in the presence of various concentrations of ethoxzolamide for 3 days, after which virus load in the culture supernatants was determined. Data are from four independent experiments, with the graph line connecting mean values. $TCID_{50}$, median tissue culture infectious dose; N.D., not detected. See also Figure 7.

**Figure S8. Harnessing of transfer learning for automated lead compound optimization**

Inspired by the idea of reinforcement learning, LIGHTHOUSE could act as an “environment” for further refinement of lead compounds. A series of new virtual compounds is generated by the “Agent” and submitted to LIGHTHOUSE (“Environment”). Output scores for the target protein (“Reward”) are then harnessed for the next generation of virtual compounds. Iteration of these cycles should allow the “Agent” to learn what are much improved compounds for the protein of interest.

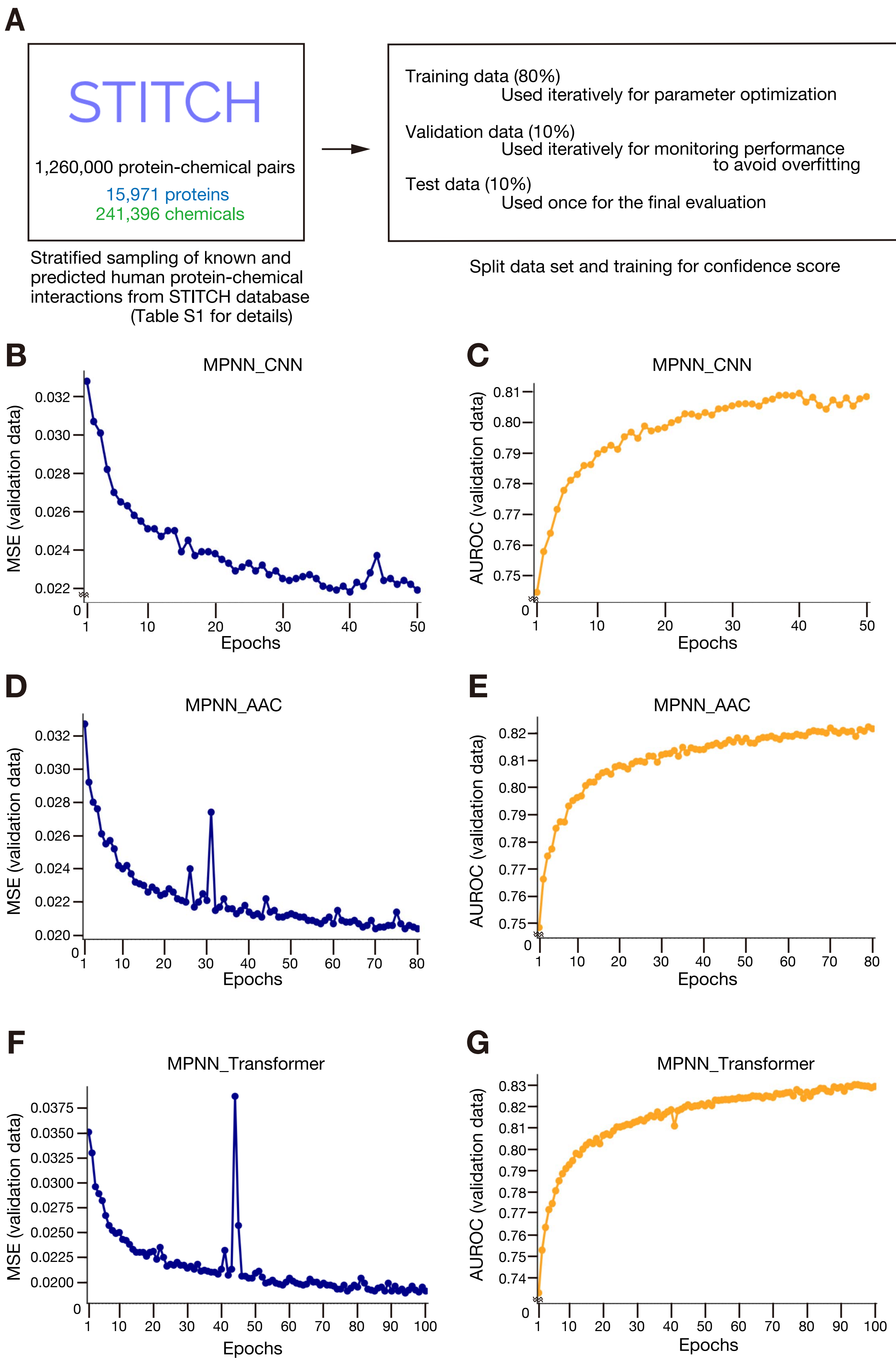


**Shimizu et al., Figure S1**

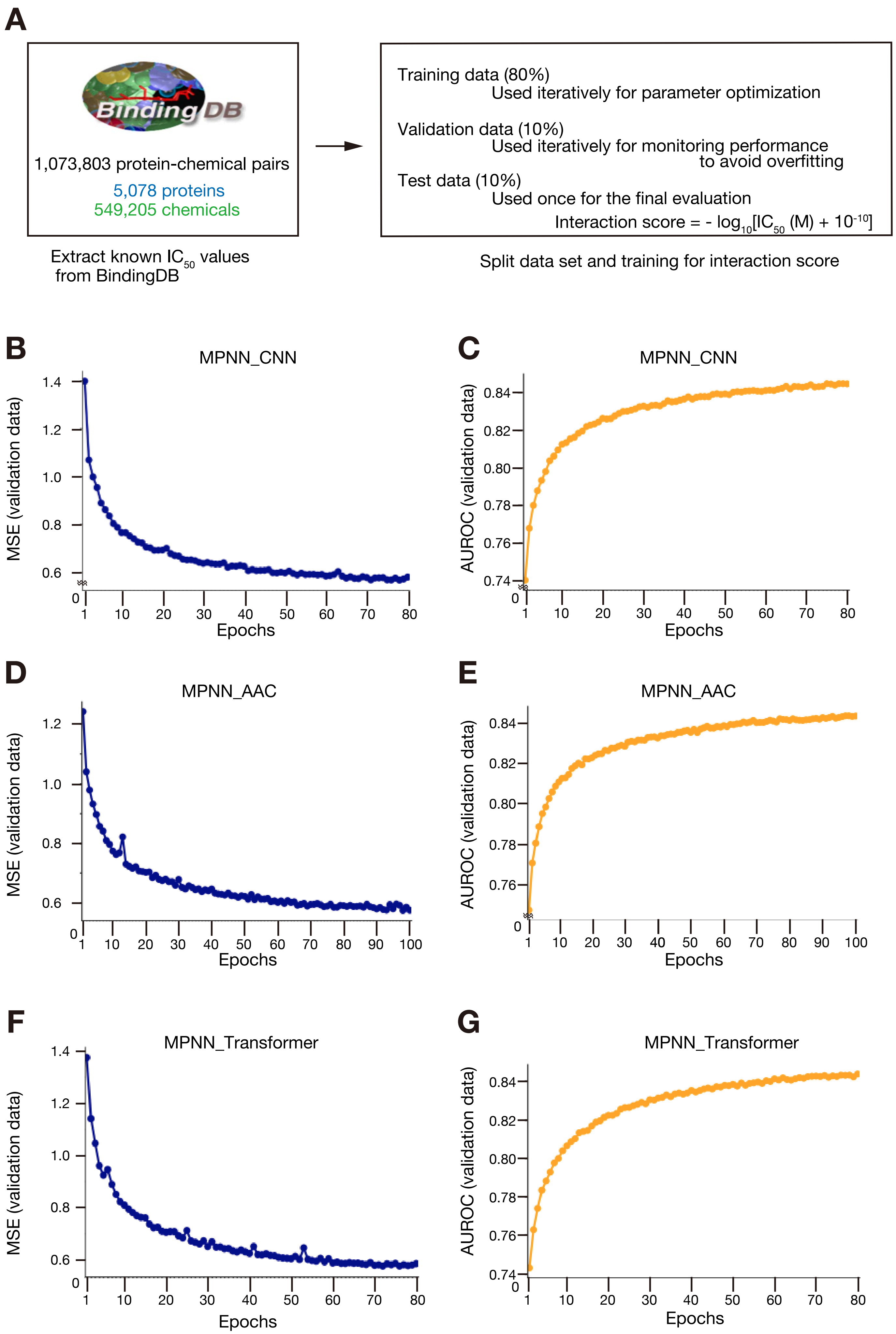


**Shimizu et al., Figure S2**

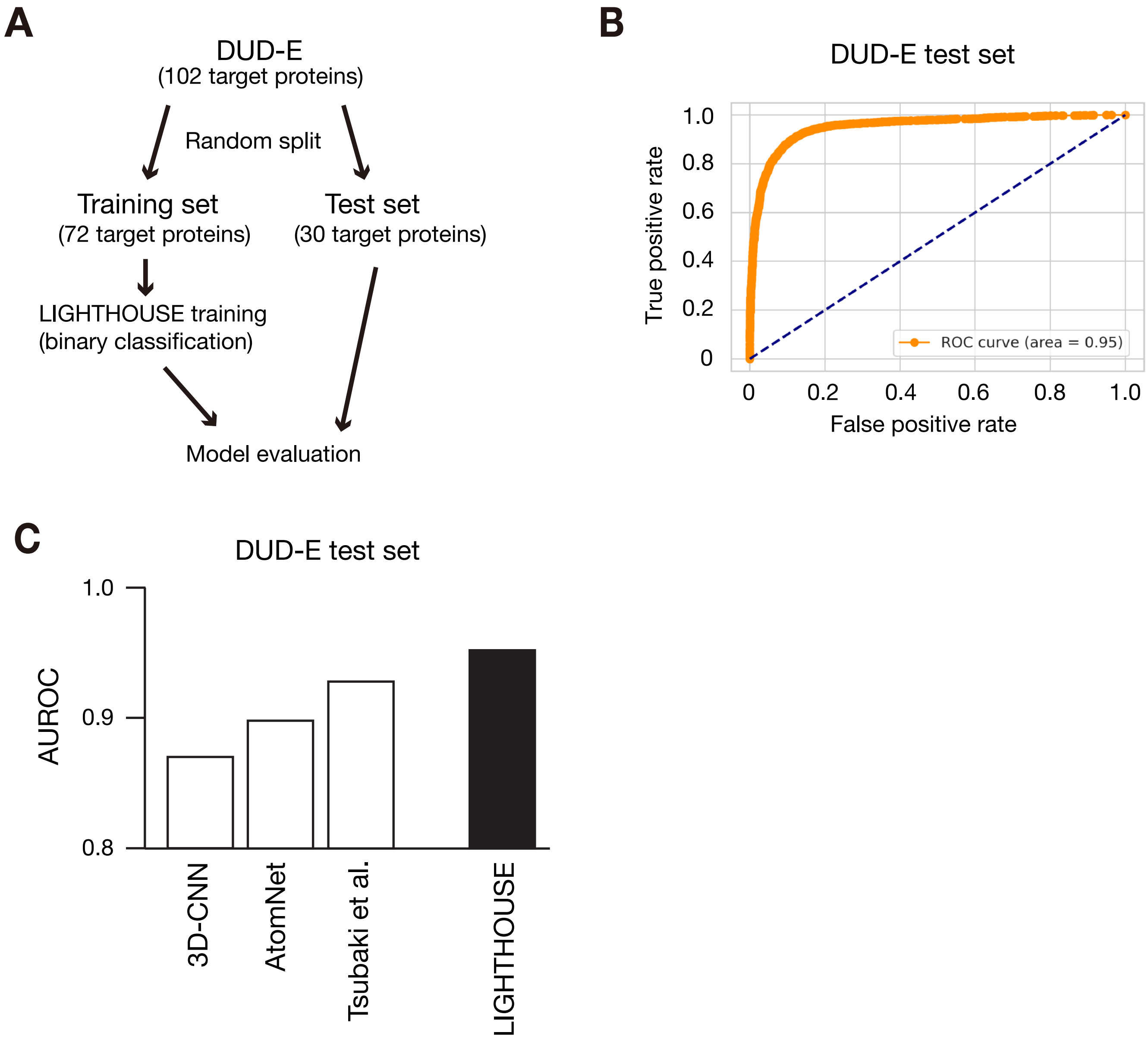


**Shimizu et al., Figure S3**

BindingDB test set (107294 CPIs)

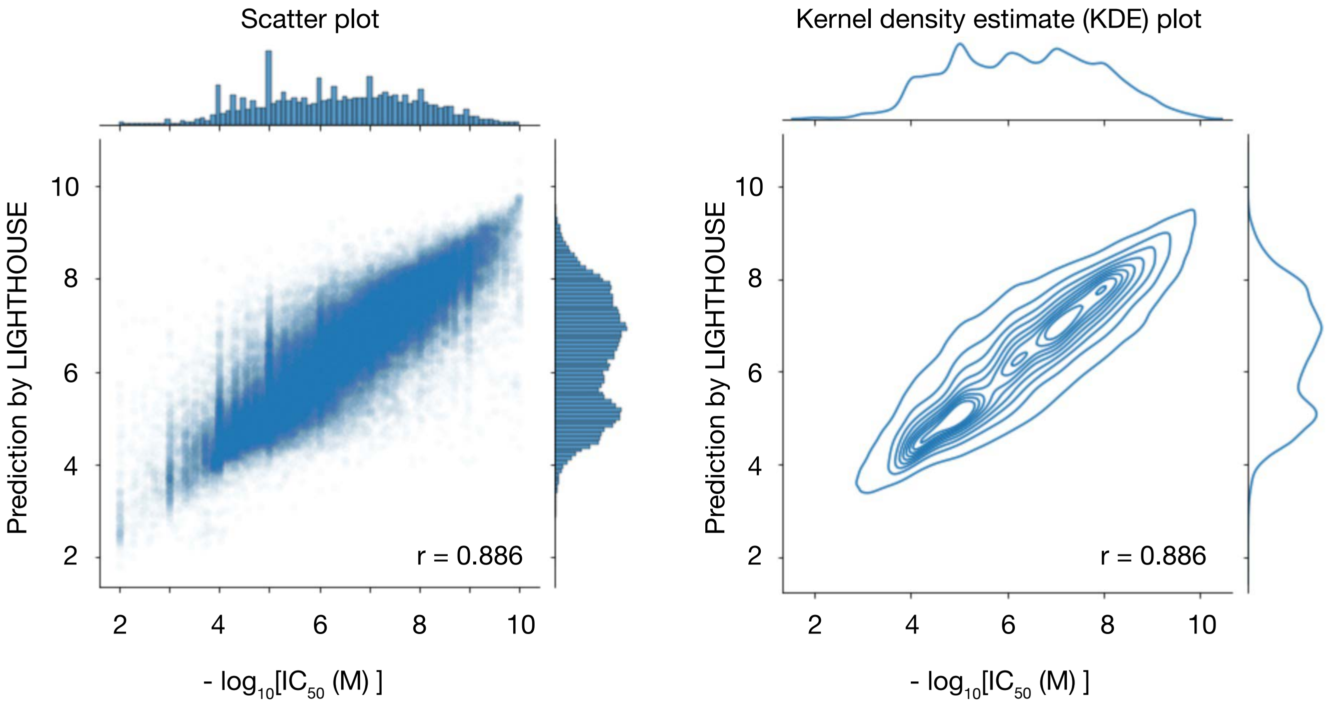


**Shimizu et al., Figure S4**

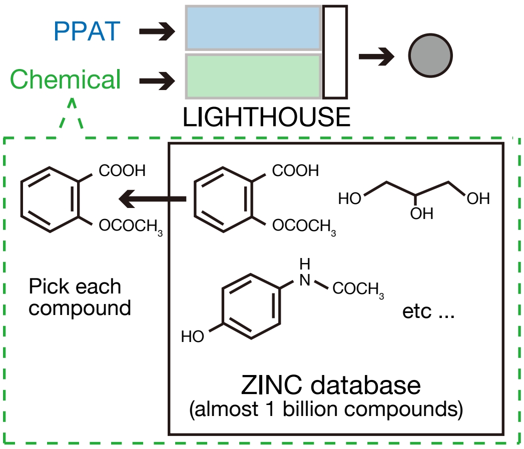


**Shimizu et al., Figure S5**

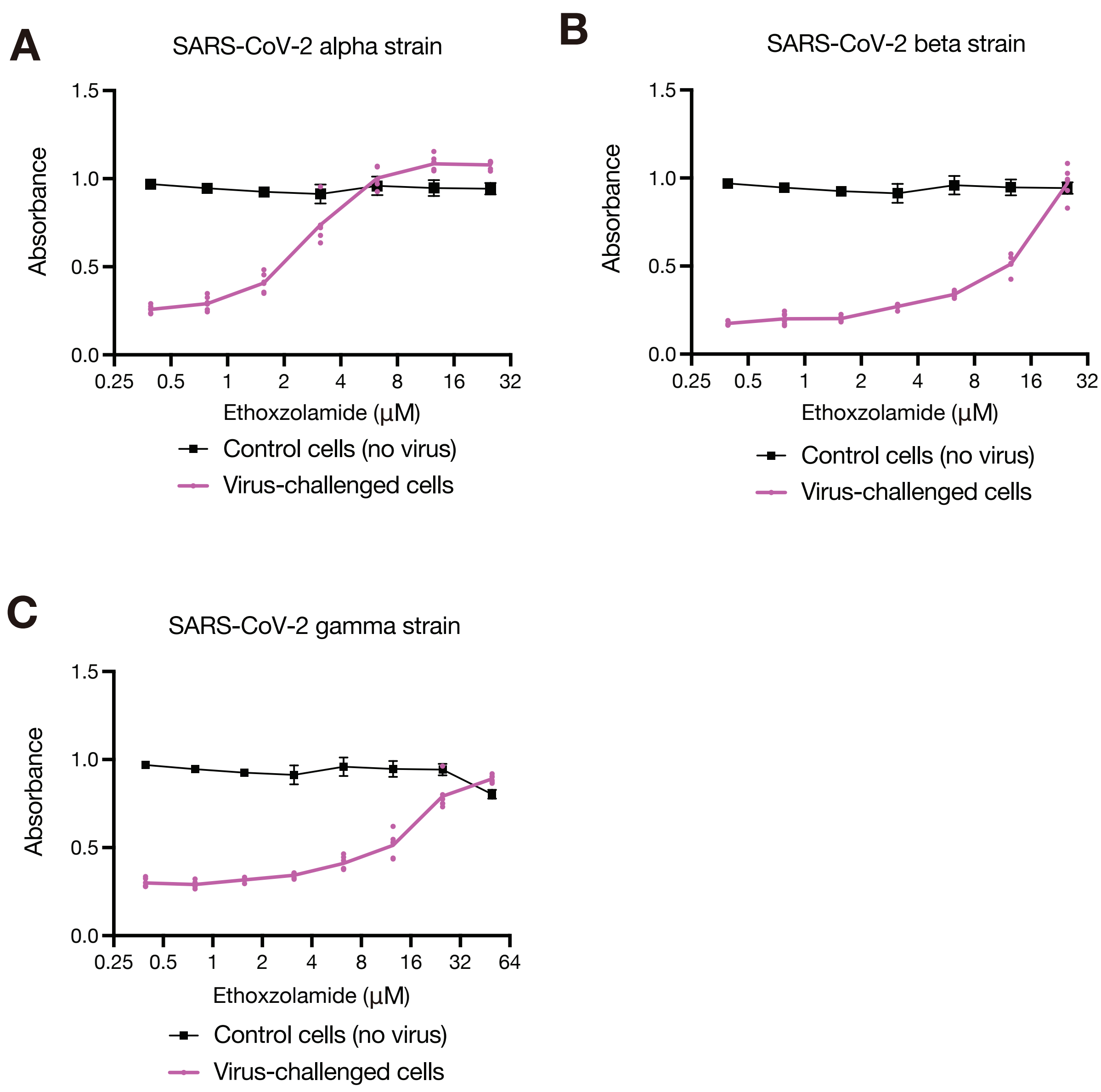


**Shimizu et al., Figure S6**

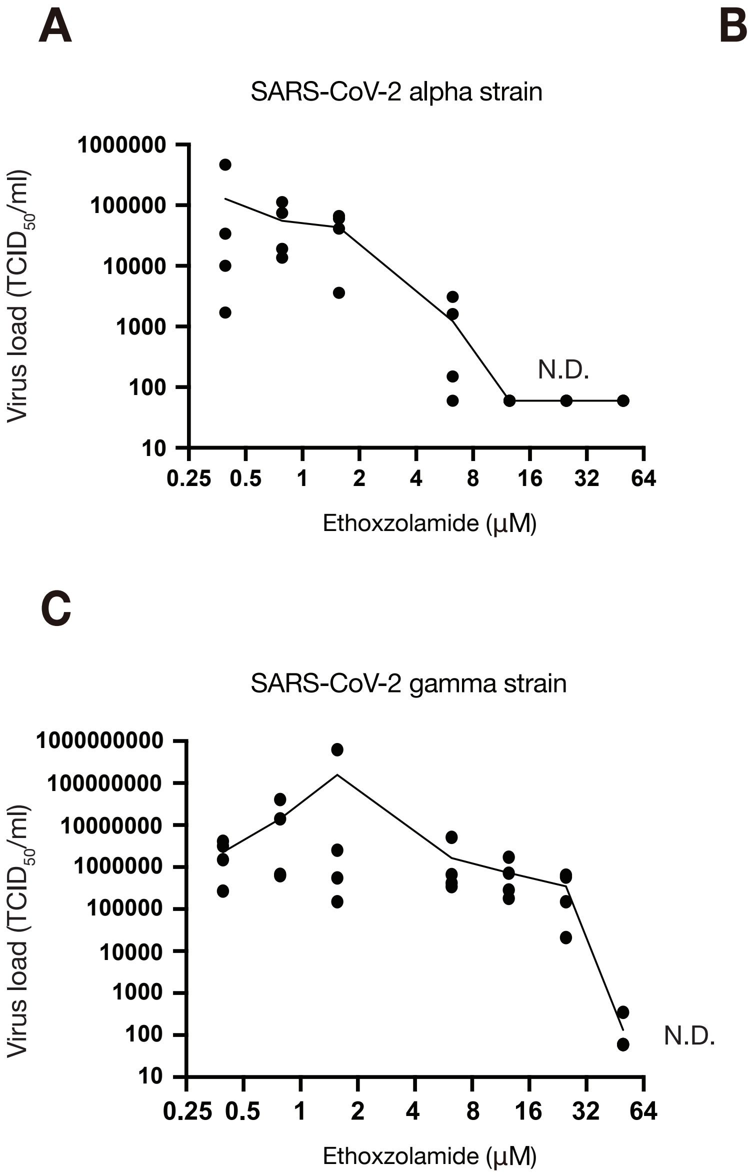


**Shimizu et al., Figure S7**

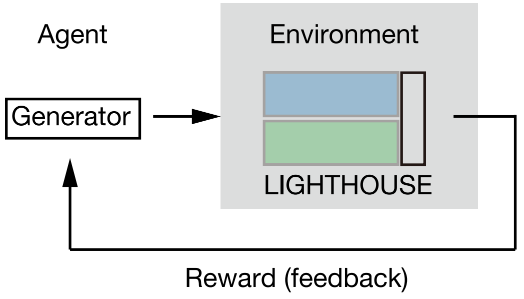


**Shimizu et al., Figure S8**
